# Viral RNA persistence in sheep tissues following acute Crimean-Congo hemorrhagic fever virus infection

**DOI:** 10.1101/2025.11.13.688338

**Authors:** Hongzhao Li, Estella Moffat, Katherine Handel, Michelle Nebroski, Daniel Sullivan, Carissa Embury-Hyatt, Oliver Lung, Bradley S Pickering

**Affiliations:** National Centre for Foreign Animal Disease, Canadian Food Inspection Agency, Winnipeg, Manitoba, Canada; Department of Medical Microbiology and Infectious Diseases, College of Medicine, Faculty of Health Sciences, University of Manitoba, Winnipeg, Manitoba, Canada

**Keywords:** Crimean-Congo hemorrhagic fever virus, Kosovo Hoti, viral RNA persistence, animal, sheep, tissue, replication, reactivation

## Abstract

Crimean-Congo hemorrhagic fever orthonairovirus (CCHFV) is a tick-borne RNA virus that can cause a severe hemorrhagic disease in humans. In animals, CCHFV infection is known to produce a transient viremia followed by host recovery and thus a short window in which animals can serve as potential intermediate hosts. Here we report that, in domestic sheep experimentally infected with CCHFV Kosovo Hoti, viral whole genome sequences were recovered from tissues long after the clearance of viremia. While viral RNA persistence in tissues was largely characterized by a lack of continuous and extensive viral replication, the sporadic presence of replication products (RNA transcripts and proteins) was detected. These findings suggest that the biology of CCHFV in animals goes far beyond an acute infection. A possibility of viral reactivation from a dormant state, akin to that observed in other RNA viruses, warrants further investigation.

## Introduction

CCHFV is a risk group 4 virus and one of the top priority pathogens listed by the World Health Organization in urgent need for research and development [1-4]. Infection in humans can lead to a range of disease outcomes, including a severe, often fatal, hemorrhagic fever disease, characterized by vascular dysfunction, hemorrhagic manifestations, multi-organ failure, shock and death [1, 5-7]. Case fatality rates are high, ranging typically from 5% to 30% and reportedly up to 80% in some outbreaks [1, 8-10]. However, no internationally licensed vaccines or therapeutics are currently available against CCHFV. Maintained and transmitted primarily by ticks from the genus *Hyalomma*, the virus is endemic in countries across Asia, Europe and Africa, with emerging signs of geographic expansion and rising global incidence of infection [11].

CCHFV belongs to the genus *Orthonairovirus*, family *Nairoviridae*, order *Bunyavirales* [12]. The virion contains a tri-segmented, single-stranded, negative-sense RNA genome (genomic vRNA) [13]. The small (S), medium (M) and large (L) genomic segments encode the nucleoprotein (N protein), glycoproteins and RNA-dependent RNA polymerase (RdRp), respectively. Genomic vRNA, N protein and RdRp together form the genomic ribonucleoprotein complexes (RNPs) inside an envelope with embedded glycoproteins. During viral replication, genomic vRNA is transcribed by RdRp to generate messenger RNA (mRNA) and genomic complementary RNA (genomic cRNA or antigenomic RNA), both of which are positive-sense. mRNA is translated into viral proteins, whereas genomic cRNA, assembling into antigenomic RNPs with N protein and RdRp, subsequently serves as a template for genomic vRNA replication [13]. In each genomic vRNA or cRNA segment, intra-strand base-pairing between the complementary terminal non-coding regions contributes to a panhandle-like secondary structure and a circular genomic appearance [13].

CCHFV circulates in a tick-vertebrate-tick cycle [1, 14]. Ticks are the primary vector and long-term reservoir of the virus. A variety of wild and domestic animals can be productively infected and develop a viremia while not showing obvious signs of disease. These animals can act as intermediate hosts amplifying and transmitting the virus. Due to their short viremic period (a few days), however, animals are not considered long-term viral carriers. While earlier experimental infection studies together with serological surveys have established a prominent role of animals in CCHFV maintenance and transmission, the investigations have been limited to the assessment of viremia and antibodies. The biology of CCHFV infection in animals remains poorly understood in many aspects. Among major unknowns are viral dissemination and persistence potentially occurring in tissues of infected animals [14]. New insights gained by addressing such gaps in CCHFV knowledge should serve to better guide professional safety practice and public health measures.

In this report, we present evidence that viral whole genome RNA sequences persisted in tissues of CCHFV-infected sheep weeks after the host recovery from an acute viremia, which was accompanied by the presence of replication-related viral RNA transcripts and protein. These data raise the possibility that the course and impact of CCHFV infection in animals may last longer than previously recognized. Consistent with our findings in CCHFV, similar phenomena of viral RNA persistence after the clearance of acute infection have been reported in other non-retroviral RNA viruses such as Measles, Zika, Ebola, Marburg and severe acute respiratory syndrome coronavirus 2 (SARS-CoV-2) viruses [15, 16]. In this context, we also discuss unanswered questions and suggest future directions toward a better understanding of the emerging biology of CCHFV RNA persistence.

## Materials and Methods

### Animal work, ethics and biosafety approval statement

Experimental infection of domestic sheep with CCHFV Kosovo Hoti and related animal work ethics were described previously [11]. Experiments involving infectious CCHFV were all conducted in the containment level 4 laboratory at the National Centre for Foreign Animal Diseases (NCFAD), Canadian Food Inspection Agency, following the institutional standard operating procedures.

### Amplification of viral RNA sequences from sheep tissue homogenates by reverse transcription-nested polymerase chain reaction (RT-nested PCR)

CCHFV cell culture, sheep tissue homogenization, sample inactivation and RNA extraction were performed as previously described [11, 17, 18]. RT-nested PCR strategies were developed to improve amplification specificity and yield, as outlined in Appendix Figures 1 and 2. These typically included two or three rounds of PCR reactions following an RT reaction. The PCR reactions mostly involved a mix of multiple forward primers or/and a mix of multiple reverse primers. The RT reaction was conducted using Maxima H Minus First Strand cDNA Synthesis Kit (ThermoFisher Scientific, K1652) or ProtoScript II First Strand cDNA Synthesis Kit (New England Biolabs, E6560L). PCR rounds were performed using the LongAmp Taq PCR Kit (New England Biolabs, E5200S). In some cases, the RT and first round PCR were run as a one-step RT-PCR using SuperScript III One-Step RT-PCR System with Platinum Taq DNA polymerase (ThermoFisher Scientific, 12574018). Details of RT and PCR formulations, conditions and primer sequences are provided in Appendix Materials and Methods and Appendix Tables 1 and 2.

**Figure 1.**
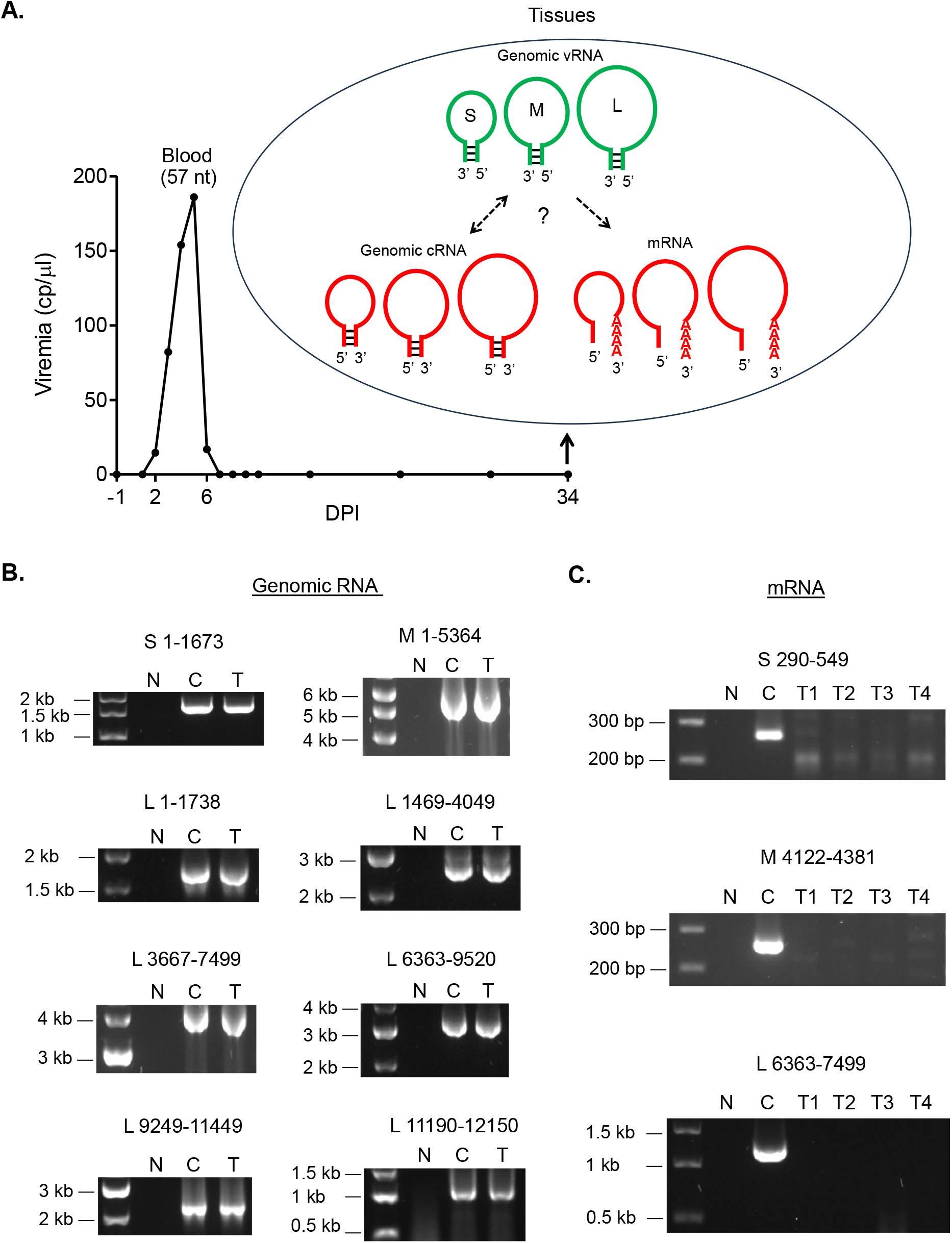
Post-acute CCHFV RNA persistence in sheep tissues. **A**. Illustration of potential viral persistence in tissues in the context of viremia dynamics. Line graph shows time course of mean viral load in whole blood of CCHFV-infected sheep, measured as viral RNA copy (cp) number/µl blood by RT-qPCR targeting a 57-nt (nucleotide) sequence of the S segment. An acute viremia was detected largely between 2 – 6 days post-infection (DPI). The 57-nt sequence was also detected in several types of tissues collected on 34 DPI, with the highest levels in lymph nodes. This raises questions concerning the format and state in which potential viral persistence may exist and prompted investigations into whether full-length CCHFV genomic segments persisted and whether viral replication-related RNA transcripts were present in these tissues. Genomic vRNA, genomic viral RNA (negative sense). Genomic cRNA, anti-genomic complementary viral RNA (positive sense). mRNA, viral messenger RNA (positive sense). **B**. Recovery of full-length CCHFV genomic sequences from sheep tissues. These were amplified by RT-nested PCR (illustrated in Appendix Figures 1 and 2), as a single amplicon for the S and M segments and as overlapping amplicons for the L segment due to large size. Amplicon name above agarose gel of PCR products indicates target region on the genomic segment relative to the reference sequence (GenBank accession number DQ133507.1 for S, EU037902.1 for M or EU044832.1 for L). N, negative control without RNA template. C, cell culture of CCHFV. T, tissue (lymph node) of CCHFV-infected sheep representing results from multiple sheep. PCR products were purified and sequenced to recover whole genome sequences. **C**. Undetectable levels of viral mRNA in sheep tissues. Purified Poly(A) RNA was analyzed by RT-nested PCR (illustrated in Appendix Figure 7). Gel images are shown as in B. T1 – T4, lymph nodes from different sheep. PCR product bands from cell culture were confirmed by sequencing to be of expected CCHFV sequences.

**Figure 2.**
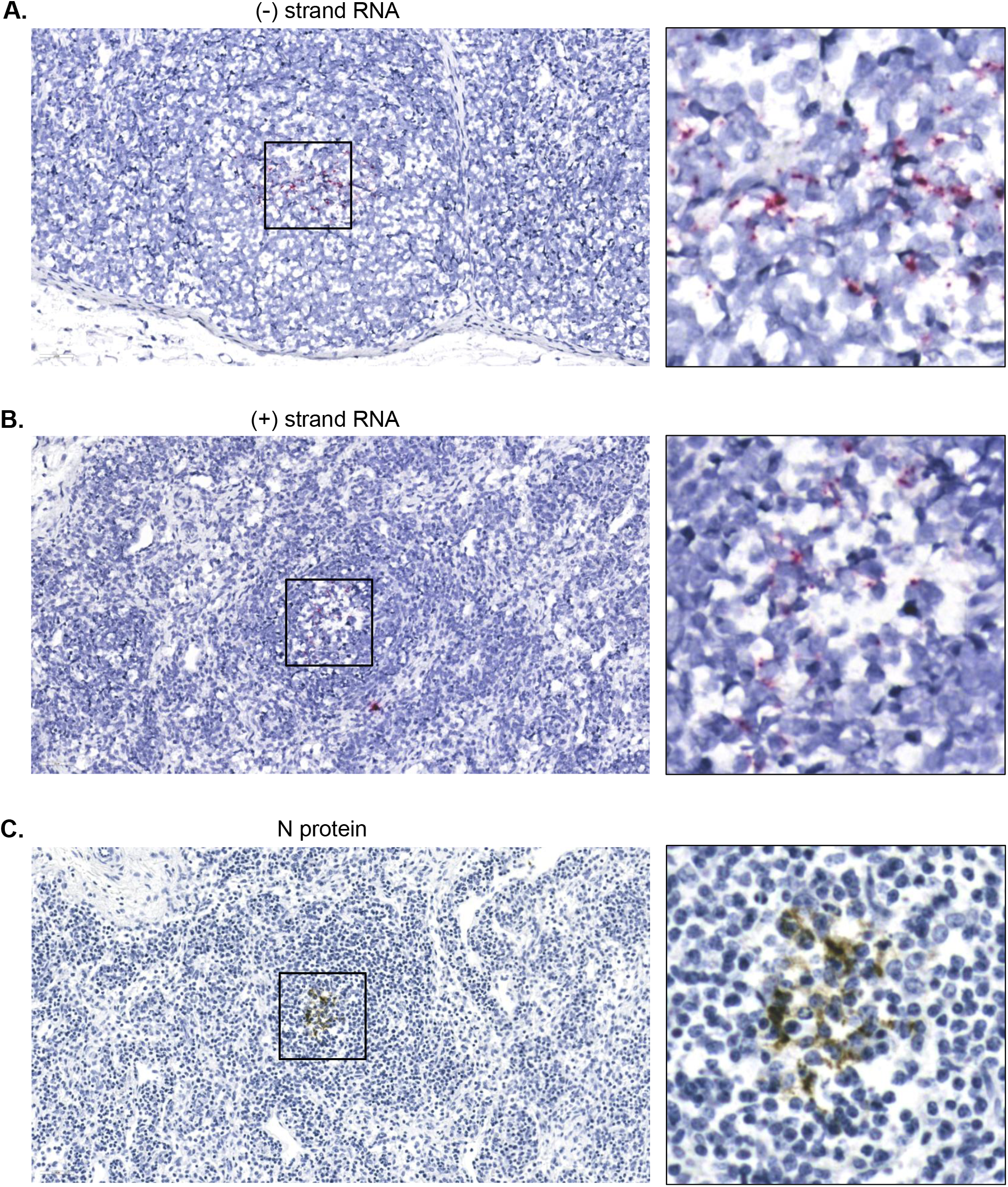
Visualization of CCHFV RNA and protein localized in sheep tissues. In situ hybridization (ISH) targeting negative-sense (A) or positive-sense (B) CCHFV RNA sequences encoded by the M segment and immunohistochemistry (IHC) targeting CCHFV N protein (C) were performed in lymph node and spleen tissues from different sheep. Results were summarized in Appendix Table 3. Microscopic images representing positive staining are shown. They were taken at 40× magnification and scaled down in this figure to 22% relative to original size. Enlarged illustration of area highlighted with square is displayed on the right in each panel.

### Next Generation Sequencing (NGS) of CCHFV RNAs from sheep tissues

RT-nested PCR amplicons were gel-purified using the MinElute Gel Extraction Kit (Qiagen, 28606), eluted with 10 – 20 µl water. The purified amplicons were either directly used or cloned into plasmids for sequencing using the TOPO TA Cloning Kit for Sequencing, with One Shot TOP10 Chemically Competent E. coli (ThermoFisher Scientific, K457540). Amplicons were sequenced based on the Oxford Nanopore Technology (ONT), using the Native Barcoding Kit 96 (Oxford Nanopore Technologies, SQK-NBD114.96), NEBNext Ultra II End repair/dA-Tailing Module (New England Biolabs, E7546), Blunt/TA Ligase Master Mix (New England Biolabs, M0367) and NEBNext Quick Ligation Module (New England Biolabs, E6056). The concentration of the amplicons was determined using a Qubit Fluorometer and a Qubit 1X dsDNA HS Assay Kit (ThermoFisher Scientific, Q33231). 200 fmol of DNA was used for NGS following the manufacturer’s protocol (Oxford Nanopore Technologies). SpotON Flow Cells (FLO-MIN114) with R10.4.1 chemistry were used to run 50 fmol of a final pooled Library on a GridION. Plasmids were sequenced using ONT or Illumina Technology. For ONT sequencing, the Rapid PCR Barcoding Kit (Oxford Nanopore Technologies, SQK-RPB114.24) and LongAmp Taq 2× Master Mix (New England Biolabs, M0287) were used. Concentration of the plasmids was determined using the Qubit method as above. 5 ng plasmid DNA was used for NGS following the manufacturer’s protocol (Oxford Nanopore Technologies) with 28 cycles of PCR. SpotON Flow Cells (FLO-MIN114) with R10.4.1 chemistry were used to run 50 fmol of a final pooled Library on a GridION. For Illumina sequencing, the concentration of the plasmids was determined as above. Then the plasmids were diluted to the appropriate concentration and libraries were prepared using the Nextera XT DNA Library Prep Kit (Illumina, FC-131-1096) and Nextera XT Index Kit v2 (Illumina, FC-131-2004), following the Illumina protocol. Libraries were cleaned up using Beckman Coulter AMpure XP Beads and the library pool was run on the Agilent TapeStation 4200 to determine the average basepair (bp) size. The library pool concentration was determined by the Qubit Fluorometer. The library pool was denatured following the Illumina Protocol “MiSeq System Denature and Dilute Libraries Guide”. A PhiX Control v3 (Illumina, FC-110-3001) was added to the final library pool at a spike in of 1% volume: volume and then the final library was loaded into a MiSeq V2 Reagent Kit (Illumina, MS-102-2002) with a final loading concentration of 9 pM and run on an Illumina MiSeq System.

### Sequencing data analysis

For Illumina sequencing runs, initial analysis was performed by running all samples through the CFIA-NCFAD/nf-villumina (https://github.com/CFIA-NCFAD/nf-villumina, v2.0.1) Nextflow [19] workflow for quality control. First, nf-villumina removed Illumina PhiX Sequencing Control V3 reads using BBDuk (v38.96) [20], followed by adapter removal and quality filtering with fastp [21]. QC filtered reads were mapped to the CCHFV Kosovo Hoti reference sequence in Geneious Prime (v2025.0.2) (https://www.geneious.com) using the Minimap2 (v.2.24) [22] assembler to generate a 75% majority consensus sequence with a minimum 10× coverage threshold. The final consensus sequences were aligned to the reference sequence with MAFFT (v7.490) [23] in Geneious Prime to determine pairwise identity.

For ONT sequencing runs, initial analysis was performed by re-basecalling the data with Dorado (https://github.com/nanoporetech/dorado, v0.9) and then performing quality trimming with Porechop_ABI (v0.5.0) [24]. In cases where large amounts of reads were produced, trimmed reads were subsampled down to 30,000 reads using seqtk (https://github.com/lh3/seqtk, v1.3-r106). Trimmed and subsampled reads were mapped against the designated CCHFV segment reference sequence in Geneious Prime (v2025.0.2) based on the target amplicon regions using Minimap2 (v2.24) [22]. Amplicon regions were determined from the final target amplicon lengths from RT-nested PCRs and 75% majority consensus sequences with a minimum 30× coverage threshold were extracted from these regions. Final consensus sequences were aligned back to the reference sequence in Geneious Prime using MAFFT (v7.490) [23] to determine pairwise identity.

### Immunohistochemistry (IHC) and in situ hybridization (ISH)

Tissue blocks were sectioned at 5 µm and stained with haematoxylin and eosin (HE) for histopathologic examination. For IHC, paraffin tissue sections were quenched for 10 minutes in aqueous 3% hydrogen peroxide. Epitopes were then retrieved using proteinase K for 10 minutes then rinsed. The primary antibody applied to the sections was a rabbit polyclonal Anti-Nucleoprotein Crimean-Congo hemorrhagic fever virus antibody (Abcam, ab190657) and was used at a 1:2,000 dilution overnight at 4 °C. Staining was then visualized using the Dako EnVision+/HRP system (anti-rabbit) (Agilent, K400311-2) and Liquid DAB+ (Agilent, K346811-2). The sections were then counter stained with Gill’s haematoxylin. For ISH, 5 µm paraffin-embedded formalin-fixed tissue sections were cut, air dried and melted onto charged slides in a 60 °C oven. The slides were then cleared and hydrated in xylene and 100% ethanol and air dried. The sections were quenched for 10 minutes in aqueous H_2_O_2_, boiled in target retrieval solution for 15 minutes, rinsed in 100% ethanol and air dried again. A final pre-treatment of protease plus enzyme for 30 minutes at 40 °C was applied. The RNAscope Probe, V-CCHFV-M-env-sense (Advanced Cell Diagnostics, 497341) or V-CCHFV-M-env (Advanced Cell Diagnostics, 479781) was applied and incubated at 40 °C for 2 hours. Hybridization amplification steps (AMP 1-6) were applied to the slides for the recommended times and temperatures as per the manual for the RNAscope 2.5 HD Reagent Kit - RED (Advanced Cell Diagnostics, 322350). The signal was visualized by the chromogen Fast Red. The sections were then counter stained with Gill’s 1 hematoxylin, dried and coverslipped. Microscopic images were acquired using the Aperio LVI real-time digital pathology system and Aperio ImageScope slide viewing software version 12.4.3.

## Results

### Signs of potential CCHFV persistence was observed in sheep tissues in the context of host recovery from an acute viremia

We recently conducted an experimental infection in domestic sheep with CCHFV Kosovo Hoti strain, where all animals developed a viremia based on the detection of a 57-nucleotide (nt) sequence of the S segment by RT-qPCR [11, 17, 18]. For a simple illustration of the viremic time range, we present the mean viral loads (copy number/µl blood) of these animals over time (Figure 1A). The viral loads were positive only within a time window of 2 – 6 days post-infection (DPI) and remained negative for weeks throughout later time points, demonstrating a brief and transient nature of the viremia. In this context, potential viral dissemination and long-term persistence in tissues were evaluated using samples collected on 34 DPI, the end point of the experimental infection study (Figure 1A). In our preliminary test, the same RT-qPCR method generated positive detections in several tissue types, with the highest levels of viral RNA present in lymph nodes, from which we also obtained near full-length sequences of the S segment [11]. The findings prompted further investigation into the characteristics of CCHFV RNA in these tissues, including whether the viral RNA retains the complete genome segments with a full coding capacity. It proved technically challenging, however, to detect from these tissues full-length genome segments or large portions of them, especially for the M and L segments. Conventional RT-PCR methods consistently generated negative results (data not shown). We reason that this might have at least in part contributed to the lack of data in the literature on CCHFV genomic RNA in tissues of animals after recovery from acute infections.

### Technical advances led to the recovery of CCHFV whole genome RNA sequences persisting in tissues

We developed RT-nested PCR strategies to address challenges in the amplification of long templates with low copy numbers from a complex tissue background (illustrated in Appendix Figures 1 and 2). Among these was an “internal nesting” to overcome the outcompeting of long, intended amplicons by short, non-specific amplicons. We established numerous RT-nested PCR conditions and primer sets to effectively amplify different genomic RNA sequences (detailed in Materials and Methods, Appendix Materials and Methods and Appendix Table 1). As a result, CCHFV whole genome RNA sequences were successfully amplified from lymph node tissues, including the full-length S and M segments each as a single amplicon and L segment as overlapping amplicons (Figure 1B). These were sequenced by NGS methods (Appendix Figures 3-5). We then performed multiple alignment analysis comparing the tissue-derived viral sequences against the Kosovo Hoti reference sequences from GenBank and the sequences of the original stock virus used to infect animals. A few scattered mutations were identified (Appendix Figures 3-5). However, none of them was consistently found in more than one animal. The absence of recurrent mutations suggests that CCHFV RNA persistence in tissues may be an inherent capability encoded in the innate viral sequences, independent of acquiring new functions through certain genomic mutations.

### Viral RNA persistence in tissues was not associated with the continuous and extensive presence of infectious virions or viral mRNA

We next asked whether CCHFV RNA persistence in tissues involves active viral replication. Some RNA viruses, namely hepatitis C virus (HCV) and Borna disease virus (BDV), are known to persist by continuous viral replication while evading immune clearance [15]. In a variety of other RNA viruses, however, viral RNA can persist without the concurrent production of infectious virions, except during occasional, late reactivation events as reported in some of these viruses [15]. To assess similar possibilities in CCHFV-infected sheep, we made several attempts of virus isolation, with our standard method of applying supernatants of tissue homogenates to SW-13 cell culture [11], but failed to recover infectious virus. While these results suggest a lack of infectious virions in the tissues of the current state, we could not certainly rule out the potential of tissues with CCHFV RNA persistence in supporting viral reactivation, which may be better addressed by more suitable and sensitive methods (discussed further below).

Alternatively, we sought to examine RNA transcripts indicative of viral replication. These are the positive-sense replicative intermediates consisting of genomic cRNA and mRNA. To distinguish them from negative-sense genomic vRNA sequences in total tissue RNA extracts, we designed strand-specific RT-nested PCR methods. Following an RT reaction with a positive sense-targeting primer, RNA was digested, RT enzyme inactivated and the resulting cDNA affinity-purified prior to being used as the template for PCR, to prevent any non-specific RT reaction from being primed by PCR primers recognizing negative RNA strands (Appendix Figure 6A). These methods generated positive detections of S segment sequences with cell culture and tissue-derived templates (Appendix Figure 6B). Interestingly, however, results remained positive in control tests with no RT primer added (Appendix Figure 6B). This suggests a false priming of the RT reaction independent of any primer addition and consequently impaired strand-specific detection. False priming was observed in previous assays intended to target the strand polarity of replicative intermediates in various viruses including hantaan virus, a bunyavirus [25]. Consistent with this possibility, positive results were similarly produced using in vitro-transcribed negative-sense S segment RNA as the template in the presence of either an RT primer targeting the negative sense or an RT primer targeting the positive sense (Appendix Figure 6C), or even in the absence of any RT primer (Appendix Figure 6D). For all the positive detections without an RT primer, the possibility of DNA template contamination was excluded as they required the addition of RT enzyme (Appendix Figure 6D). These results therefore confirm an impaired strand-specificity for the detection of positive-sense S segment transcripts. Similar findings were obtained for M segment transcripts with cell culture and tissue-derived templates (Appendix Figure 6E) and for L segment transcripts with cell culture-derived templates (Appendix Figure 6F). Underlying mechanisms of false RT priming may be explained by the snapped-back priming model proposed previously for other viruses [25], or additionally by nonspecific priming by a cellular sequence in the context of total RNA extracts from tissues (Appendix Figure 6G). Together, these data add to the literature for strand-specific detection of viral replicative intermediates by including CCHFV as a new target virus with lessons learned and nevertheless provide additional evidence of the presence of persisting CCHFV RNA sequences using alternative assays.

To improve the detection specificity of replication markers, we then chose to focus on targeting viral mRNA. Poly(A) RNA in total RNA extracts was pull-down purified using oligo(dT)-coupled magnetic beads, with all other RNA washed off, followed by RT-nested PCR analysis (Appendix Figure 7). mRNA sequences derived from the S, M and L segments, respectively, were detected from CCHFV cell culture but not from tissues (Figure 1C). Although we could not rule out the possible presence of viral mRNA in tissues at low levels beyond the detection limit of the method used, these data support that CCHFV RNA persistence in these tissues was largely in a quiescent state without extensive viral replication as seen in cell culture.

### Visualization of viral RNA persistence in tissues revealed the presence of replication products

To evaluate directly in tissues the presence and distribution patterns of potential CCHFV RNA and protein, ISH targeting negative-sense or positive-sense viral RNA (derived from the M segment) and IHC targeting viral protein (the N protein) were developed (Appendix Figures 8-10) and performed in lymph node and spleen tissues (Appendix Table 3). Both negative-sense and positive-sense viral RNA populations were observed in multiple tested tissues (Appendix Table 3 and Figure 2A and B). Viral protein was detected in one of the lymph node tissues (Appendix Table 3 and Figure 2C). Without widespread presence, the viral RNA and protein demonstrated a sporadic localized distribution (Appendix Figures 11-13). The presence of viral positive-sense RNA and protein indicates the occurrence of recent replication activities in tissues. We reason that the positive-sense RNA was predominantly, if not exclusively, genomic cRNA, since viral mRNA was not found at detectable levels (Figure 1C). Viral RNA persistence was therefore largely constituted by genomic vRNA and genomic cRNA. The lack of viral mRNA may be explained by a more rapid turnover of mRNA whereas genomic cRNA was maintained longer, or by a possibility that its production was inhibited or restricted to minimal levels while genomic cRNA transcription was allowed to occur at higher levels. With these further illustrated, we provide a hypothetical model in which we summarize the current findings and propose possible mechanisms supporting the establishment and maintenance of CCHFV RNA persistence in tissues (Appendix Figure 14). In this context, we highlight a need for future investigation into the potential of viral reactivation harbored in such persistence.

## Discussion

### The form of CCHFV RNA persistence

The detection of a CCHFV sequence from sheep tissues by RT-qPCR long after acute infection (Figure 1A) may lead to the assumption that this represents remnants of infection in the form of fragmented or degraded viral RNA. However, such nonfunctional remnants would be unlikely to persist for those several weeks, as RNA is highly susceptible to rapid degradation and clearance in the absence of cellular protective mechanisms. It has been observed in several RNA viruses that long-term dormant viral forms possess the potential to resume mRNA transcription, protein synthesis and infectious virus production, attesting to the persistence of full-length genomic RNA [15, 26-33]. Consistent with this, we recovered CCHFV full-length S and M segments from sheep tissues. We were only able to obtain the whole sequence of the L segment using overlapping PCR amplicons. No method was available from the literature or our own custom development to amplify the L segment as a single amplicon, even with high concentration template CCHFV RNA extracted from cell culture supernatants. Thus, we could not definitely confirm the full-length nature of the L segment sequences. Nevertheless, the recovery of full-length S and M segments strongly suggests the presence of a functional biological mechanism(s) required to maintain their long-term persistence in intact forms.

In addition to the possession of full-length genome segments, CCHFV RNA persistence in tissues exhibited several other features: the presence of both viral genomic vRNA and cRNA, lack of detectable viral mRNA, lack of culturable infectious virions, rare occasion of detectable viral protein, as well as a sporadic localized distribution of viral RNA and protein. These illustrate a largely dormant state without continuous and extensive production of infectious virions, but do not exclude the possibility of genomic replication (vRNA and cRNA) independent of a concurrent virion production or the possibility of sporadic viral replication. It is noteworthy that mRNA generally has a rapid turnover [34]. A subtle and brief viral mRNA transcription could go undetected. The lack of recoverable virus may also be attributed to an inadequate viral isolation method [15] or/and an inactive state of viral RNA persistence at the time of tissue harvest, outside narrow windows of potential virion production (to be further elaborated below).

### Mechanisms of CCHFV RNA persistence

It is currently unknown how CCHFV RNA persistence was initiated and maintained. We integrate findings in CCHFV with what is known in other RNA viruses and propose possible mechanisms (outlines illustrated in Appendix Figure 14). We characterized CCHFV RNA persistence mostly in lymph node tissues with highest levels of viral RNA by RT-qPCR. Among cell types found in lymph nodes, dendritic cells, macrophages and endothelial cells are known target cells of CCHFV infection. The role of lymphocytes (B cells and T cells) including memory B and T cells is less clear in permitting CCHFV replication. Which of these cell types are involved in viral RNA persistence remains to be determined. CCHFV replicates in the cytoplasm, which is therefore a likely site for viral RNA to persist in host cells. Viral RNA may be protected as RNPs or by association with membrane structures [15]. Some non-retroviral RNA viruses such as BDV [35] and SARS-CoV-2 [36] can generate a cDNA version of their genomic sequences through the activity of host endogenous retroviral elements. This did not appear to be the case for CCHFV as the detection of viral sequences by RT-PCR was RT enzyme-dependent, excluding the presence of DNA template (Appendix Figure 6D).

CCHFV RNA persistence may be mediated by viral and cellular mechanisms that downregulate viral replication to prevent lethal damage to the host cell and the elimination of the persisting viral RNA or the host cell by immune responses. Mutations in measles virus promote intercellular transfer of viral RNPs without continued production of infectious virions [15]. This mechanism does not seem to be used by CCHFV since no recurrent mutation was identified in the persisting genomic RNA sequences. Interferon (IFN) responses are a known antiviral mechanism that restricts CCHFV replication [37]. Susceptibility to lethal infection has been established in IFN-deficient knockout animals [1, 2]. IFN-stimulated genes encode proteins with antiviral activities such as RNA degradation and inhibition of virion production [38]. On the other hand, previous studies suggest that the CCHFV genome encodes IFN-antagonizing viral functions [37]. It is tempting to speculate that, depending on the type and physiologic state of host cells, a balance achieved between the host antiviral responses and the viral counter mechanisms may facilitate the suppression of mRNA transcription and virion production as well as the initiation and establishment of viral RNA persistence. This may occur in an infected cell following a period of viral replication, or right after viral entry (Appendix Figure 14). To maintain long-term persistence, genomic vRNA and cRNA may be carried in long-lived cells such as memory B cells or T cells or replicated and segregated into the daughter cells following the division of proliferating cells. They may also be cell-to-cell transferred as RNPs to neighboring cells as in measles virus [39].

It is a key question to address whether persistent CCHFV RNA sequences can react to changes in immune control or cellular physiologic state and begin replicating infectious virions, which can fuel new cycles of viral RNA persistence and potentially cause late viral transmissions. This possibility was hinted by anti-Gn and N protein antibody waves at late time points of the experimental infection study [11]. Viral reactivation from dormancy has been documented in several other non-retroviral RNA viruses in response to treatments that weaken immune control [15, 26-30]. Factors associated with viral reactivation have been more widely studied in HIV and DNA viruses, most of which can potentially affect immune control and might therefore be relevant to non-retroviral RNA viruses including CCHFV. These factors include immunosuppression, other viral infections, physical trauma, physiological changes (e.g. fever, menstruation and hormonal fluctuations), environmental factors (e.g. sunlight exposure) and host cell changes (e.g. differentiation and gene expression) [40, 41]. Viral reactivation from CCHFV RNA persistence may be sporadic and short-lived due to the subsequent activation of host memory immune responses. The evaluation of reactivatable CCHFV replication will require methods that maintain the viability of host cells. Tissue homogenization-based viral isolation should be unsuitable by failing to conserve the intact cellular machinery that can support potential viral replication. Candidate factors for viral reactivation such as immunosuppression (through drug treatment with cyclophosphamide) can be tested in animals or in vitro co-culture systems with freshly isolated tissues or cells carrying viral RNA persistence [29, 30, 42-44].

### Consequence and impact of CCHFV RNA persistence

Viral RNA persistence in other RNA viruses has been linked to recurring, chronic or progressive post-viral disease or syndromes in the hosts and late transmissions of infectious virus that can lead to new outbreaks [15, 16]. Of veterinary relevance, although CCHFV does not cause prominent clinical signs in animals such as sheep, long-term health effects of viral RNA persistence should only be excluded by controlled experimental studies, as these effects could potentially go unrecognized or regarded as non-specific variations in health status among the animal population. CCHFV RNA persisting in lymph nodes may serve as a source of recurring antigenic stimulations, as suggested by the late antibody waves [11]. Persistent immune activation has been implicated in immune exhaustion and increased re-infection by SARS-CoV-2 [45]. However, continued antigenic boost in lymphoid tissues is believed to facilitate long-term immunity against measles virus [46, 47]. These different possibilities warrant follow-up investigations in CCHFV-infected sheep.

The possibility of viral reactivation from CCHFV RNA persistence remains to be determined or ruled out by further experimental studies. Before then, it represents an additional, previously unappreciated source of potential viral transmission. Due to the high consequence of CCHFV infection, this information should be made available to professionals at high risk such as slaughterhouse workers and veterinarians through cautionary public health education and measures.

Epidemiological surveillance of CCHFV infections in animals commonly depends on serological tests for CCHFV antibodies whereas the use of more sensitive and specific molecular methods such as RT-qPCR for viral RNA is limited by the short and transient nature of viremia. However, viral RNA persistence in tissues, which can be collected during livestock slaughtering, provides an extended window for detection and thus a novel opportunity to evaluate CCHFV RNA prevalence in animal populations.

### Study limitation and future directions

The major limitation of the current study was only one time point of tissue collection. Our recent experimental infection study, which provided the tissue materials, was a pilot test initially designed to characterize CCHFV Kosovo Hoti infection in sheep, without any knowledge of viral dissemination and persistence in tissues, and thus included only one late time point of animal sacrifice for tissue collection following the 3Rs animal use ethics [11]. If CCHFV RNA persistence is dynamic, rather than static, involving cycles of biological events (Appendix Figure 14), the same single time point, although in multiple animals, will likely have limited coverage of all its aspects. An extended future investigation with serial tissue sampling time points will facilitate the in-depth characterization of viral RNA persistence. This will provide opportunities to address the temporal and spatial sequence of viral dissemination through tissues, the duration of viral RNA persistence and the molecular, cellular and immunological mechanisms contributing to the initiation, establishment and maintenance of viral RNA persistence in a spatiotemporal context. Of particular importance, potential viral reactivation can be evaluated using animals or tissues from different time points, in combination with inducing treatments such as with an immunosuppressive drug.

## Appendix

A single PDF file including supplementary materials and methods, tables, figures and references.

## Acknowledgements

This work was supported by a Canadian Safety and Security Program grant (CSSP-2018-CP-2341) and funding from Canadian Food Inspection Agency.

## Declaration of interest statement

The authors declare that they have no competing interests.

## Notes

### Competing Interest Statement

The authors have declared no competing interest.

