## Supplemental Materials for "Viral RNA persistence in sheep tissues following acute Crimean-Congo hemorrhagic fever virus infection"

### Table of contents

Appendix Materials and Methods……………………..……………..……….……………………………......2

Appendix Table 1. RT-PCR conditions and primer sequences for amplifying CCHFV genomic RNA sequences.4

Appendix Table 2. RT-PCR conditions and primer sequences for targeting the strand polarity of CCHFV RNA transcripts……………………..………………………………………………………………………....5

Appendix Table 3. Visualization of CCHFV RNA and protein in tissues……….…………………….………..6

Appendix Figure 1. RT-nested PCR to amplify full-length CCHFV S or M genomic RNA from sheep tissues….7

Appendix Figure 2. RT-nested PCR to amplify CCHFV L genomic RNA sequences from sheep tissues………...8

Appendix Figure 3. Sequence of CCHFV S segment from sheep tissues……………...……………………………9

Appendix Figure 4. Sequence of CCHFV M segment from sheep tissues………………..……..…………...…..10

Appendix Figure 5. Sequence of CCHFV L segment from sheep tissues.…..…………..…………………...…..11

Appendix Figure 6. RT-nested PCR targeting strand polarity of CCHFV RNA transcripts in total RNA extracts from sheep tissues…………………………………………………………………………………………...12

Appendix Figure 10. Establishment of immunohistochemistry (IHC) for CCHFV N protein…………………...16

Appendix Figure 11. Image of ISH for negative-sense CCHFV RNA (M segment) displaying a larger area of tissue section…………………………………………………………………………………………...17

Appendix Figure 12. Image of ISH for positive-sense CCHFV RNA (M segment) displaying a larger area of tissue section……………………………………………………………………………………………………18

Appendix Figure 13. Image of IHC for CCHFV N protein displaying a larger area of tissue section…………...19

Appendix Figure 14. Hypothetical model of CCHFV RNA persistence in tissues………………………………20

References……………………………………………………………..………………...…………21

### Appendix Materials and Methods

#### Amplification of viral genomic RNA sequences by RT-nested PCR

The RT-nested PCR strategies to amplify CCHFV genomic sequences are outlined in Appendix Figures 1 and 2. While the following describes the common RT-PCR formulations and PCR conditions, amplicon-specific variations in reaction conditions and primer sequences can be found in Appendix Table 1. For RT reaction using the Maxima H Minus First Strand cDNA Synthesis Kit (ThermoFisher Scientific, K1652), a denaturation/annealing reaction (15 µl) containing 8 µl water, 1 µl 10 µM or 100 µM RT primer, 5 µl RNA extraction and 1 µl 10 mM dNTP mix was performed with the thermal program: Denaturation at 65 °C for 5 minutes and annealing at 4°C for 5 minutes. After addition of 4 µl 5× RT Buffer and 1 µl Maxima H Minus Enzyme Mix, the final RT reaction (20 µl) was run with the thermal program: RT at 50 °C or 65 °C for 30 minutes, RT enzyme inactivation at 85 °C for 5 minutes and incubation at 4 °C for 5 minutes. For RT reaction using the ProtoScript II First Strand cDNA Synthesis Kit (New England Biolabs, E6560L), a denaturation/annealing reaction (8 µl) containing 2 µl water, 1 µl 10 µM RT primer and 5 µl RNA extraction was performed with the same thermal program as above with the Maxima H Minus RT. After addition of 10 µl 2× ProtoScript II Reaction Mix and 2 µl 10× ProtoScript II Enzyme Mix, the final RT reaction (20 µl) was run with the thermal program: RT at 42 °C for 60 minutes, RT enzyme inactivation at 85 °C for 5 minutes and incubation at 4 °C for 5 minutes.

For PCR rounds using the LongAmp Taq PCR Kit (New England Biolabs, E5200S), the first round PCR reaction (20 µl) consisted of 8 µl water, 4 µl 5× LongAmp Taq Reaction Buffer, 0.6 µl 10 mM dNTP mix, 0.8 µl 10 µM (total primer concentration) forward primer or primer mix, 0.8 µl 10 µM (total primer concentration) reverse primer or primer mix, 0.8 µl LongAmp Taq and 5 µl template (RT products). The second or third round PCR reaction had the same formulation except that the volumes of water and template (previous round PCR products) were 12 µl and 1 µl, respectively. The thermal program for LongAmp Taq PCR was: pre-denaturation at 94 °C for 30 seconds, 40 amplification cycles (denaturation at 94 °C for 10 seconds, annealing at 65 °C for 1 minute and extension at 65 °C for a customized time depending on the amplicon size, largely 50 seconds/kb), final extension at 65 °C for 10 minutes and cooling down at 4 °C for 5 minutes. The final PCR products were visualized on a 0.8% agarose gel with the molecular size marker 1 kb DNA Ladder (New England BioLabs, N3232S). DNA bands were purified and sequenced as below.

For one-step RT-PCR (combining RT and the first round of nested PCRs) using the SuperScript III One-Step RT-PCR System with Platinum Taq DNA polymerase (ThermoFisher Scientific, 12574018), the reaction (50 µl) consisted of 16 µl water, 25 µl 2× Reaction Mix, 2 µl SuperScript III RT/Platinum Taq Mix, 1 µl 10 mM forward primer mix, 1 µl 10 mM reverse primer mix and 5 µl RNA template. The thermal program was: RT at 52.5 °C for 30 minutes, pre-denaturation at 94 °C for 2 minutes, 40 amplification cycles (denaturation at 94 °C for 15 seconds, annealing at 50.5 °C for 30 seconds and extension at 68 °C for 1 minute 40 seconds), final extension at 68 °C for 5 minutes and cooling down at 4 °C for 5 minutes. 1 µl of the resulting RT-PCR products was used as the template in a subsequent 20 µl LongAmp Taq PCR reaction.

#### RT-nested PCR targeting the strand polarity of CCHFV RNA transcripts

The strand polarity-specific RT-nested PCR strategy is outlined in Appendix Figure 6A. Regarding the common reaction formulations and conditions, RT reactions were performed similarly to the above using the Maxima H Minus First Strand cDNA Synthesis Kit (ThermoFisher Scientific, K1652) except that the Maxima H Minus Enzyme Mix (containing RNase inhibitor RiboLock) was replaced by Maxima H Minus Reverse Transcriptase (ThermoFisher Scientific, EP0752, containing no RNase inhibitor) and only a 100 µM RT primer master stock was used. PCR reactions were performed using the LongAmp Taq PCR method as described above. Amplicon-specific variations in reaction conditions and primer sequences can be found in Appendix Table 2. RT reactions were each based on an RT primer designed to recognize a target viral RNA sequence of a specific strand polarity (Appendix Figure 6A). To remove RNA and RT enzyme and purify cDNA, 20 µl RT reaction was digested with 1 µl RNase H (New England BioLabs, M0297L) and 1 µl RNase A from the RNaseAlert Lab Test Kit v2

(ThermoFisher Scientific, 4479768) at 37 °C for one hour and then at room temperature overnight, followed by purification with MinElute Gel Extraction Kit (Qiagen, 28604).

**Production of the S segment RNA genome by in vitro transcription with enhanced DNA removal (S-IVT)**
The negative-sense S segment RNA genome of CCHFV Kosovo Hoti was in vitro-transcribed, treated with DNase I and column-purified as previously described [1]. This was diluted at 1:200 in H2O and further treated with the DNA-free DNA Removal Kit (ThermoFisher Scientific, AM1906), in a digestion reaction consisting of 83 µl H2O, 5 µl diluted RNA, 10 µl 10× Reaction Buffer and 2 µl rDNase with incubation at 37 °C for 30 minutes. The digestion reaction was then cleaned up using the RNeasy Mini Kit (Qiagen, 74104) with RNA elution in 30 µl H2O. The eluted RNA was further treated with restriction enzyme digestion in a reaction containing 22 µl H2O, 20 µl RNA, 5 µl 10× NEBuffer EcoRI/SspI and 3 µl EcoRI (New England Biolabs, R0101S) with incubation at 37 °C for one hour. The digestion reaction was finally cleaned up using the RNeasy Mini Kit with RNA elution in 30 µl Core elution buffer from the MagMAX CORE Nucleic Acid Purification Kit (ThermoFisher Scientific, A32700).

**Detection of viral mRNA sequences by RT-nested PCR**
mRNA was purified from total RNA extracts from tissue homogenates with Dynabead mRNA Purification Kit (ThermoFisher Scientific, 61006) by following the supplier's protocol (https://assets.thermofisher.com/TFS-Assets/LSG/manuals/MAN0015808_Dynabeads_mRNA_Purification_UG.pdf) with these customizations: 40 µl input RNA was combined with 60 µl 10 mM Tris-HCl, pH 7.5 to serve as the starting material. mRNA bound to the beads was finally eluted in two rounds: first round with 20 µl Tris-HCl, heated at 78°C for 2 minutes; second round with 5 µl, heated at 80°C for approximately 4 minutes. Each heating step was immediately followed by placing the tube on the magnet for collection of the mRNA elute separated from the beads. The elutes from both rounds were finally combined and analyzed by positive sense-targeting RT-nested PCR as detailed in Appendix Figure 7 and Appendix Table 2.

**Appendix Table 1. RT-PCR conditions and primer sequences for amplifying CCHFV genomic RNA sequences**

| Amplicon name | Amplicon type | Location of final amplicon on genomic segment | Final amplicon size (bp) | Reverse transcription (RT) and Nested PCR round 1 (one step or two steps) | | | | Nested PCR round 2 | | | Nested PCR round 3 | | |
|---|---|---|---|---|---|---|---|---|---|---|---|---|---|
| | | | | RT | Nested PCR round 1 | | RT and PCR conditions | | | | | | |
| | | | | (Primer name:<br>Primer sequence) | Forward primer or primer pool<br>(Primer name:<br>Primer sequence) | Reverse primer or primer pool<br>(Primer name:<br>Primer sequence) | | Forward primer or primer pool<br>(Primer name:<br>Primer sequence) | Reverse primer or primer pool<br>(Primer name:<br>Primer sequence) | PCR conditions | Forward primer or primer pool<br>(Primer name:<br>Primer sequence) | Reverse primer or primer pool<br>(Primer name:<br>Primer sequence) | PCR conditions |
| S 1-1673 | Full-Length S | Nucleotide positions 1 – 1673 of S segment (DQ133507.1) | 1673 | fS1:<br>TCTCAAAGAAACACGTGCCGC | | rS1:<br>TCTCAAAGATATCGTTGCCGC | RT-PCR:<br>SuperScript III One-Step RT-PCR System with Platinum Taq DNA Polymerase, with extension time: 1m40s | fS2:<br>TCTCAAAGAAACACGTGCCGCTTACG<br><br>fS3:<br>TCTCAAAGAAACACGTGCCGCTTACGCCCA<br><br>fS4:<br>TCTCAAAGAAACACGTGCCGCTTACGCCCACAGT<br><br>fS5:<br>TCTCAAAGAAACACGTGCCGCTTACGCCCACAGTGTTCTC<br><br>fS6:<br>TCTCAAAGAAACACGTGCCGCTTACGCCCACAGTGTTCTCTTGAGTGTCTG | rS2:<br>TCTCAAAGATATCGTTGCCGCACA<br><br>rS3:<br>TCTCAAAGATATCGTTGCCGCACAGCCCT<br><br>rS4:<br>TCTCAAAGATATCGTTGCCGCACAGCCCTTTAA<br><br>rS5:<br>TCTCAAAGATATCGTTGCCGCACAGCCCTTTAAGTATTTG<br><br>rS6:<br>TCTCAAAGATATCGTTGCCGCACAGCCCTTTAAGTATTTGAAATGAAAGATAG | LongAmp Taq PCR, with annealing/extension time: 2m24s | | | |
| M 1-5364 | Full-Length M (RT-PCR method 1) | Nucleotide positions 1 – 5364 of M segment (EU037902.1) | 5364 | fM1:<br>TCTCAAAGAAATACTTGCGG<br><br>fM2:<br>TCTCAAAGAAATACTTGCGGCA<br><br>fM3:<br>TCTCAAAGAAATACTTGCGGCACGT<br><br>fM4:<br>TCTCAAAGAAATACTTGCGGCACGTCAGTAC<br><br>fM5:<br>TCTCAAAGAAATACTTGCGGCACGTCAGTACGTAAGTG<br><br>fM6:<br>TCTCAAAGAAATACTTGCGGCACGTCAGTACGTAAGTGTCAACTTTGAGA | fM1:<br>TCTCAAAGAAATACTTGCGG<br><br>fM2:<br>TCTCAAAGAAATACTTGCGGCA<br><br>fM3:<br>TCTCAAAGAAATACTTGCGGCACGT<br><br>fM4:<br>TCTCAAAGAAATACTTGCGGCACGTCAGTAC<br><br>fM5:<br>TCTCAAAGAAATACTTGCGGCACGTCAGTACGTAAGTG<br><br>fM6:<br>TCTCAAAGAAATACTTGCGGCACGTCAGTACGTAAGTGTCAACTTTGAGA | rM1:<br>TCTCAAAGATATAGTGGCGG<br><br>rM2:<br>TCTCAAAGATATAGTGGCGGCACG<br><br>rM3:<br>TCTCAAAGATATAGTGGCGGCACGCAGTCAAAGG<br><br>rM4:<br>TCTCAAAGATATAGTGGCGGCACGCAGTCAAAGGTAAGGTCAGA<br><br>rM5:<br>TCTCAAAGATATAGTGGCGGCACGCAGTCAAAGGTAAGGTCAGAGCAAGCAGAG | RT:<br>ProtoScript® II First Strand cDNA Synthesis Kit<br><br>PCR:<br>LongAmp Taq PCR, with annealing/extension time: 11m8s | fM8:<br>TCTCAAAGAAATACTTGCGGCACGTCAGTACGTAAGTGTCAACTTTGAGAGAGTGGATTGTGC<br><br>fM9:<br>TCTCAAAGAAATACTTGCGGCACGTCAGTACGTAAGTGTCAACTTTGAGAGAGTGGATTGTGCATCCCGAATGCACCA | rM6:<br>TCTCAAAGATATAGTGGCGGCACGCAGTCAAAGGTAAGGTCAGAGCAAGCAGAGCTAGGGCAGT<br><br>rM7:<br>TCTCAAAGATATAGTGGCGGCACGCAGTCAAAGGTAAGGTCAGAGCAAGCAGAGCTAGGGCAGTTTCAAGC | LongAmp Taq PCR, with annealing/extension time: 11m8s | | | |
| M 1-5364 | Full-Length M (RT-PCR method 2) | Nucleotide positions 1 – 5364 of M segment (EU037902.1) | 5364 | fM1:<br>TCTCAAAGAAATACTTGCGG<br><br>fM2:<br>TCTCAAAGAAATACTTGCGGCA<br><br>fM3:<br>TCTCAAAGAAATACTTGCGGCACGT<br><br>fM4:<br>TCTCAAAGAAATACTTGCGGCACGTCAGTAC<br><br>fM5:<br>TCTCAAAGAAATACTTGCGGCACGTCAGTACGTAAGTG<br><br>fM6:<br>TCTCAAAGAAATACTTGCGGCACGTCAGTACGTAAGTGTCAACTTTGAGA | fM1:<br>TCTCAAAGAAATACTTGCGG<br><br>fM2:<br>TCTCAAAGAAATACTTGCGGCA<br><br>fM3:<br>TCTCAAAGAAATACTTGCGGCACGT<br><br>fM4:<br>TCTCAAAGAAATACTTGCGGCACGTCAGTAC<br><br>fM5:<br>TCTCAAAGAAATACTTGCGGCACGTCAGTACGTAAGTG<br><br>fM6:<br>TCTCAAAGAAATACTTGCGGCACGTCAGTACGTAAGTGTCAACTTTGAGA | rM1:<br>TCTCAAAGATATAGTGGCGG<br><br>rM2:<br>TCTCAAAGATATAGTGGCGGCACG<br><br>rM3:<br>TCTCAAAGATATAGTGGCGGCACGCAGTCAAAGG<br><br>rM4:<br>TCTCAAAGATATAGTGGCGGCACGCAGTCAAAGGTAAGGTCAGA<br><br>rM5:<br>TCTCAAAGATATAGTGGCGGCACGCAGTCAAAGGTAAGGTCAGAGCAAGCAGAG | RT:<br>ProtoScript® II First Strand cDNA Synthesis Kit<br><br>PCR:<br>LongAmp Taq PCR, with annealing/extension time: 11m8s | fM7:<br>TCTCAAAGAAATACTTGCGGCACGTCAGTACGTAAGTGTCAACTTTGAGAGAG<br><br>fM8:<br>TCTCAAAGAAATACTTGCGGCACGTCAGTACGTAAGTGTCAACTTTGAGAGAGTGGATTGTGC | rM6:<br>TCTCAAAGATATAGTGGCGGCACGCAGTCAAAGGTAAGGTCAGAGCAAGCAGAGCTAGGGCAGT | LongAmp Taq PCR, with annealing/extension time: 11m8s | | | |
| L 1-1738 | L upstream end | Nucleotide positions 1 – 1738 of L segment (EU044832.1) | 1738 | fL0:<br>TCTCAAAGATATCAATCCCCCCTTTACCCA<br><br>fL1:<br>TCTCAAAGATATCAATCCCCCCTTTACCCACGTTG<br><br>fL2:<br>TCTCAAAGATATCAATCCCCCCTTTACCCACGTTGACACAGT<br><br>fL3:<br>TCTCAAAGATATCAATCCCCCCTTTACCCACGTTGACACAGTCAGCTCC | | L1a-R5:<br>GTCTCTTGATCTCGATCAGAAGCTG<br><br>L1a-R6:<br>TCTACTTCAGAATCTTCTTGAAGAGCATA | RT-PCR:<br>SuperScript III One-Step RT-PCR System with Platinum Taq DNA Polymerase, with extension time: 1m50s | fL4:<br>TCTCAAAGATATCAATCCCCCCTTTACCCACGTTGACACAGTCAGCTCCAGTGGTG<br><br>fL4b:<br>TCTCAAAGATATCAATCCCCCCTTTACCCACGTTGACACAGTCAGCTCCAGTGGTGGATTTTT<br><br>fL4c:<br>TCTCAAAGATATCAATCCCCCCTTTACCCACGTTGACACAGTCAGCTCCAGTGGTGGATTTTTGTCTCT<br><br>fL4d:<br>TCTCAAAGATATCAATCCCCCCTTTACCCACGTTGACACAGTCAGCTCCAGTGGTGGATTTTTGTCTCTGTGGTACC | L1a-R3:<br>CTCAAGTAAAGGTTATGTGCATAGGA<br><br>L1a-R4:<br>TGAAATATCCTCAGAGGTCATATTTA | LongAmp Taq PCR, with annealing/extension time: 2m32s | | | |
| L 1-1678 | L upstream end | Nucleotide positions 1 – 1678 of L segment (EU044832.1) | 1678 | CCHF-L1F:<br>TCTCAAAGATATCAATCCCCCC<br>(10 µM RT primer stock used) | fL0:<br>TCTCAAAGATATCAATCCCCCCTTTACCCA<br><br>fL1:<br>TCTCAAAGATATCAATCCCCCCTTTACCCACGTTG<br><br>fL2:<br>TCTCAAAGATATCAATCCCCCCTTTACCCACGTTGACACAGT<br><br>fL3:<br>TCTCAAAGATATCAATCCCCCCTTTACCCACGTTGACACAGTCAGCTCC | L1a-R5:<br>GTCTCTTGATCTCGATCAGAAGCTG<br><br>L1a-R6:<br>TCTACTTCAGAATCTTCTTGAAGAGCATA | RT:<br>Maxima H Minus First Strand cDNA Synthesis Kit, with reaction temperature: 50 °C<br><br>PCR:<br>LongAmp Taq PCR, with annealing/extension time: 2m32s | fL4:<br>TCTCAAAGATATCAATCCCCCCTTTACCCACGTTGACACAGTCAGCTCCAGTGGTG<br><br>fL4b:<br>TCTCAAAGATATCAATCCCCCCTTTACCCACGTTGACACAGTCAGCTCCAGTGGTGGATTTTT<br><br>fL4c:<br>TCTCAAAGATATCAATCCCCCCTTTACCCACGTTGACACAGTCAGCTCCAGTGGTGGATTTTTGTCTCT<br><br>fL4d:<br>TCTCAAAGATATCAATCCCCCCTTTACCCACGTTGACACAGTCAGCTCCAGTGGTGGATTTTTGTCTCTGTGGTACC | L1a-R3:<br>CTCAAGTAAAGGTTATGTGCATAGGA<br><br>L1a-R4:<br>TGAAATATCCTCAGAGGTCATATTTA | LongAmp Taq PCR, with annealing/extension time: 2m32s | fL5:<br>TCTCAAAGATATCAATCCCCCCTTTACCCACGTTGACACAGTCAGCTCCAGTGGTGGATTTTTGTCTCTGTGGTACCATGGA<br><br>fL6:<br>TCTCAAAGATATCAATCCCCCCTTTACCCACGTTGACACAGTCAGCTCCAGTGGTGGATTTTTGTCTCTGTGGTACCATGGACTTCTT<br><br>fL7:<br>TCTCAAAGATATCAATCCCCCCTTTACCCACGTTGACACAGTCAGCTCCAGTGGTGGATTTTTGTCTCTGTGGTACCATGGACTTCTTGAGGAACCTT<br><br>fL8:<br>TCTCAAAGATATCAATCCCCCCTTTACCCACGTTGACACAGTCAGCTCCAGTGGTGGATTTTTGTCTCTGTGGTACCATGGACTTCTTGAGGAACCTTGATTGGACTCA<br><br>fL9:<br>TCTCAAAGATATCAATCCCCCCTTTACCCACGTTGACACAGTCAGCTCCAGTGGTGGATTTTTGTCTCTGTGGTACCATGGACTTCTTGAGGAACCTTGATTGGACTCAGGTGATTGCTG | L1a-R1:<br>CTTCCTGAGGTGCTGATGAATTTCATAAT<br><br>L1a-R2:<br>ACATTTCCAATCTTGCCTGCACTCATTG | LongAmp Taq PCR, with annealing/extension time: 2m32s |
| L 1469-4049 | L internal (RT-PCR method 1) | Nucleotide positions 1469 – 4049 of L segment (EU044832.1) | 2581 | CCHF-L1F:<br>TCTCAAAGATATCAATCCCCCC<br>(10 µM RT primer stock used) | L1b-F1:<br>AATCTGGACTTTGACTGTCTAAGTGTGCA<br><br>L1b-F2:<br>GACTATAATAACATGATAGAAAGTATGTC | L-R5:<br>ACTGTTTATAACTTCTAGACACCTATACTGCTGCA<br><br>L-R5b :<br>TCAATGTCTTGCAATGTCTTTTCATTTAC<br><br>L-R5c :<br>TGAATGCATCTTCTAGCAAAGTCATGGAGTGA | RT:<br>Maxima H Minus First Strand cDNA Synthesis Kit, with reaction temperature: 50 °C<br><br>PCR:<br>LongAmp Taq PCR, with annealing/extension time: 3m45s | L1b-F3:<br>CAAATTGGATAACACTGATGTTGAATT<br><br>L1b-F4:<br>CAACCACAGGGAGATAGCTGATCTCAA | L-R5d :<br>ACCTACCCTCAAGAGCAAAAGTGAGTTCC<br><br>L-R4:<br>CTTCATAAGGTGTGCCAATGTACCTCTTCAT<br><br>L-R4b:<br>GATCAGAGAAACTAAGGTGTCGATAATTC | LongAmp Taq PCR, with annealing/extension time: 3m45s | | | |
| L 1469-4049 | L internal (RT-PCR method 2) | Nucleotide positions 1469 – 4049 of L segment (EU044832.1) | 2581 | L1b-F1:<br>AATCTGGACTTTGACTGTCTAAGTGTGCA<br><br>L1b-F2:<br>GACTATAATAACATGATAGAAAGTATGTC | | L-R5:<br>ACTGTTTATAACTTCTAGACACCTATACTGCTGCA<br><br>L-R5b :<br>TCAATGTCTTGCAATGTCTTTTCATTTAC<br><br>L-R5c :<br>TGAATGCATCTTCTAGCAAAGTCATGGAGTGA | RT-PCR:<br>SuperScript III One-Step RT-PCR System with Platinum Taq DNA Polymerase, with extension time: 3m18s | L1b-F3:<br>CAAATTGGATAACACTGATGTTGAATT<br><br>L1b-F4:<br>CAACCACAGGGAGATAGCTGATCTCAA | L-R5d :<br>ACCTACCCTCAAGAGCAAAAGTGAGTTCC<br><br>L-R4:<br>CTTCATAAGGTGTGCCAATGTACCTCTTCAT<br><br>L-R4b:<br>GATCAGAGAAACTAAGGTGTCGATAATTC | LongAmp Taq PCR, with annealing/extension time: 3m45s | | | |
| L 3667-7499 | L internal | Nucleotide positions 3667 – 7499 of L segment (EU044832.1) | 3833 | L-F3:<br>AATGAATGCTTAACAGATCTAGACAGGCTTGTTAG<br>(100 µM RT primer stock used) | L-F3:<br>AATGAATGCTTAACAGATCTAGACAGGCTTGTTAG | CCHF-L1R:<br>TTGGCACTATCTTTCATTTGAC | RT:<br>Maxima H Minus First Strand cDNA Synthesis Kit, with reaction temperature: 50 °C<br><br>PCR:<br>LongAmp Taq PCR, with annealing/extension time: 5m | L-F4:<br>ATGTCAGGCATTGATTGCATAAAATATCCCACAGGACAGCTTATCAC | CCHF-L1R3:<br>GCATGTTCAACATGTACTGTTGT | LongAmp Taq PCR, with annealing/extension time: 5m | L-F5:<br>CTTATGCAAGAGGGGTGATTTCAAATTCACTGAAAAACCTCTC | CCHF-L1R4:<br>ACCCTGACCCATATGGTTGTAACTGTT | LongAmp Taq PCR, with annealing/extension time: 5m |
| L 6363-9520 | L internal | Nucleotide positions 6363 – 9520 of L segment (EU044832.1) | 3158 | CCHF-L2F-Hoti:<br>GAAGAGCTGTATGACATAAGGC<br>(10 µM RT primer stock used) | CCHF-L2F-Hoti2:<br>GAGCACCCCAGTGTCAGCTTCAC | L-R10:<br>TTCATCAAGTTACCTTTCAGCATGTAGAAAAC | RT:<br>Maxima H Minus First Strand cDNA Synthesis Kit, with reaction temperature: 65 °C<br><br>PCR:<br>LongAmp Taq PCR, with annealing/extension time: 5m | CCHF-L2F-Hoti3:<br>TCAGCTCCTTGGTGCTTACACTACAAAGCAC | L-R9:<br>GTTGGCTTTGCTTGTTGTATCATCCTGCTGAGCA | LongAmp Taq PCR, with annealing/extension time: 5m | CCHF-L2F-Hoti4:<br>AGACACCCAGAGATACTTGACTGTGG | L-R8:<br>AAAGTGAACACTGAGGGGAAAAACTGCATTAT | LongAmp Taq PCR, with annealing/extension time: 5m |
| L 7441-9370 | L internal | Nucleotide positions 7441 – 9370 of L segment (EU044832.1) | 1930 | L-F3:<br>AATGAATGCTTAACAGATCTAGACAGGCTTGTTAG<br>(100 µM RT primer stock used) | CCHF-L2F-Hoti:<br>GAAGAGCTGTATGACATAAGGC<br><br>CCHF-L2F-Hoti2:<br>GAGCACCCCAGTGTCAGCTTCAC<br><br>CCHF-L2F-Hoti3:<br>TCAGCTCCTTGGTGCTTACACTACAAAGCAC<br><br>CCHF-L2F-Hoti4:<br>AGACACCCAGAGATACTTGACTGTGG | L-R8:<br>AAAGTGAACACTGAGGGGAAAAACTGCATTAT<br><br>L-R9:<br>GTTGGCTTTGCTTGTTGTATCATCCTGCTGAGCA<br><br>L-R10:<br>TTCATCAAGTTACCTTTCAGCATGTAGAAAAC | RT:<br>Maxima H Minus First Strand cDNA Synthesis Kit, with reaction temperature: 50 °C<br><br>PCR:<br>LongAmp Taq PCR, with annealing/extension time: 4m | CCHF-L2F-Hoti5:<br>GAGGACTGTAAGCTCACTTTGCTTGATCT<br><br>CCHF-L2F-Hoti6:<br>TCAGTTTCCAAGCTCTTAGTCTGTTTGTATC<br><br>CCHF-L2F-Hoti7:<br>GTTACCCTATTCACATCCAATGG<br><br>CCHF-L2F-Hoti8:<br>ACTGACGAAAAGCTACTACATCA<br><br>CCHF-L2F-Hoti9:<br>ACAAAGATAGTTTTTGCTAAAAT | L-R13:<br>TTAAAGCGATCTTTAAGTAAGTTTGA<br><br>L-R14:<br>TCAACTTTGGTAGGTTTGACTTC<br><br>L-R15:<br>GAAATTGCTGATTCATCCAATAAATAGCCC | LongAmp Taq PCR, with annealing/extension time: 4m | Amp3F1:<br>CAACCTATATAAGCAAAGGACTCATGGC<br><br>Amp3F2:<br>GGAAATGACACAGTTAAGTTTTTAGT<br><br>Amp3F3:<br>TGTTGGCAGATAATTTAGACAGTTGGGA<br><br>Amp3F4:<br>GGAGGTGTAGAGCAATATAGTGAAGA<br><br>Amp3F5:<br>ATACCTGCTGGTAGCATTAAGAAGATCTTAAATGT | L-R16:<br>ATGACAACTGCAGGATTGTTGGATAGTCG<br><br>L-R17:<br>TGTTCAGTTGCCGAGAACAGTTTGGTTTTAACAAT | LongAmp Taq PCR, with annealing/extension time: 4m |
| L 9249-11449 | L internal | Nucleotide positions 9249 – 11449 of L segment (EU044832.1) | 2201 | L-F7:<br>AAGTACACTCACTGGTTGGACACCTTTCT<br>(100 µM RT primer stock used) | L-F7:<br>AAGTACACTCACTGGTTGGACACCTTTCT | CCHF-L2R:<br>TCTCAAAGAAATCGTTCCCCCCAC<br><br>CCHF-L2R2:<br>TCTCAAAGAAATCGTTCCCCCCACACCC<br><br>CCHF-L2R3:<br>TCTCAAAGAAATCGTTCCCCCCACACCCTTTAA<br><br>CCHF-L2R4:<br>TCTCAAAGAAATCGTTCCCCCCACACCCTTTAAATAATGAAACTG<br><br>L-R11:<br>TCATCCAAGCTTATTTGCTCTTGACACTGCTC<br><br>L-R12:<br>TCAATCTGAATCCCAGTTAAAGCCAAGGTTTCCTG<br><br>L-R18:<br>GTTCAATTTGTGTCACGGTACTAGGTTTG | RT:<br>Maxima H Minus First Strand cDNA Synthesis Kit, with reaction temperature: 50 °C<br><br>PCR:<br>LongAmp Taq PCR, with annealing/extension time: 4m | L-F8:<br>ATACAGAGTCATCTCAAAGTCTTTGAAAATGA<br><br>L-F9:<br>CTTCAGTTCTCCGATCCAAACAGACAGAAACT | L-R19:<br>TGCCTGTTAACATCAGAATTAAATCTTCTACTGCTCTAAC<br><br>L-R20:<br>CCTAGCTCATTGTTTGCACGAACA<br><br>L-R21:<br>ACCAAGACAATATTAACCCTCAGTGACACATC | LongAmp Taq PCR, with annealing/extension time: 4m | | | |
| L 11190-12150 | L downstream end | Nucleotide positions 11190 – 12150 of L segment (EU044832.1) | 961 | L-F7:<br>AAGTACACTCACTGGTTGGACACCTTTCT<br>(100 µM RT primer stock used) | L4bF1:<br>AAAACTACATTGAAGATATAGGCTT<br><br>L4bF2:<br>AACAATGTGCTTGTTAACATTGACAG<br><br>L4bF3:<br>AACACAGTGAAAAGTGAACTTTTGTGGAA<br><br>L4bF4:<br>ATTTACGTTAGACACTAATGTAAGCAC | CCHF-L2R:<br>TCTCAAAGAAATCGTTCCCCCCAC<br><br>CCHF-L2R2:<br>TCTCAAAGAAATCGTTCCCCCCACACCC<br><br>CCHF-L2R3:<br>TCTCAAAGAAATCGTTCCCCCCACACCCTTTAA<br><br>CCHF-L2R3b:<br>TCTCAAAGAAATCGTTCCCCCCACACCCTTTAAATAATG<br><br>CCHF-L2R4:<br>TCTCAAAGAAATCGTTCCCCCCACACCCTTTAAATAATGAAACTG<br><br>rL0:<br>TCTCAAAGAAATCGTTCCCCCCACACCCTT<br><br>rL1:<br>TCTCAAAGAAATCGTTCCCCCCACACCCTTTAAATAATGA | RT:<br>Maxima H Minus First Strand cDNA Synthesis Kit, with reaction temperature: 50 °C<br><br>PCR:<br>LongAmp Taq PCR, with annealing/extension time: 1m59s | L4bF5:<br>ACAAGCACCATTAAAGATGTGAGGACA<br><br>L4bF6:<br>CTGGTATCCTACGTCAGTACTGAAAC<br><br>L4bF7:<br>TGCTTGCATTCCTTCTCTACGAAGAAGT<br><br>L-F20:<br>ATGAACTTAATCAACCAGTGTAAGGCAGT | rL2:<br>TCTCAAAGAAATCGTTCCCCCCACACCCTTTAAATAATGAAACTGGTATG<br><br>CCHF-L2R5:<br>TCTCAAAGAAATCGTTCCCCCCACACCCTTTAAATAATGAAACTGGTATGAA<br><br>CCHF-L2R6:<br>TCTCAAAGAAATCGTTCCCCCCACACCCTTTAAATAATGAAACTGGTATGAATAGTAG<br><br>CCHF-L2R7:<br>TCTCAAAGAAATCGTTCCCCCCACACCCTTTAAATAATGAAACTGGTATGAATAGTAGAGAAAG | LongAmp Taq PCR, with annealing/extension time: 1m53s | | | |
| L 1-729 | L upstream end | Nucleotide positions 1 – 729 of L segment (EU044832.1) | 729 | CCHF-L1F:<br>TCTCAAAGATATCAATCCCCCC<br>(10 µM RT primer stock used) | fL0:<br>TCTCAAAGATATCAATCCCCCCTTTACCCA<br><br>fL1:<br>TCTCAAAGATATCAATCCCCCCTTTACCCACGTTG<br><br>fL2:<br>TCTCAAAGATATCAATCCCCCCTTTACCCACGTTGACACAGT<br><br>fL3:<br>TCTCAAAGATATCAATCCCCCCTTTACCCACGTTGACACAGTCAGCTCC | L-R3:<br>GATTCTGGCCTTAGAATTAAAGTGTCAGCTGATATCTCAACACA<br><br>L-R3b:<br>TAGTTTACAGTTAAACATGCAGTCCAAG<br><br>L-R3c:<br>ACCCTGCCGACTCTAGTCGGTATG | RT:<br>Maxima H Minus First Strand cDNA Synthesis Kit, with reaction temperature: 50 °C; or ProtoScript® II First Strand cDNA Synthesis Kit<br><br>PCR:<br>LongAmp Taq PCR, with annealing/extension time: 1m47s | fL4:<br>TCTCAAAGATATCAATCCCCCCTTTACCCACGTTGACACAGTCAGCTCCAGTGGTG<br><br>fL4b:<br>TCTCAAAGATATCAATCCCCCCTTTACCCACGTTGACACAGTCAGCTCCAGTGGTGGATTTTT<br><br>fL4c:<br>TCTCAAAGATATCAATCCCCCCTTTACCCACGTTGACACAGTCAGCTCCAGTGGTGGATTTTTGTCTCT<br><br>fL4d:<br>TCTCAAAGATATCAATCCCCCCTTTACCCACGTTGACACAGTCAGCTCCAGTGGTGGATTTTTGTCTCTGTGGTACC | L-R2:<br>TTCTGACTTTTATTGACCAATTTAGACAACTGAGATGC<br><br>L-R2b:<br>CTTCTGCCTCAATATCTCTGCTTGTTTCC<br><br>L-R2c:<br>TGGACAGAGTGACTTCATCCAAGCTTG | LongAmp Taq PCR, with annealing/extension time: 1m47s | | | |
| L 301-4049 | L internal | Nucleotide positions 301 – 4049 of L segment (EU044832.1) | 3749 | L-F1:<br>TTTGAGATTGTGCGGCAGCCTGGTGATGG<br><br>L-F1b:<br>GAACTGCTTCTACCACAGCATAGCTGAG<br><br>L-F1c:<br>ACCATGCCCAACAAAACAGCTCACTC | | L-R5:<br>ACTGTTTATAACTTCTAGACACCTATACTGCTGCA<br><br>L-R5b :<br>TCAATGTCTTGCAATGTCTTTTCATTTAC<br><br>L-R5c :<br>TGAATGCATCTTCTAGCAAAGTCATGGAGTGA | RT-PCR:<br>SuperScript III One-Step RT-PCR System with Platinum Taq DNA Polymerase, with extension time: 4m32s | L-F1d:<br>ATATCACAACATCAAACATCTGACCGAG<br><br>L-F1e:<br>TTGGCAGCAAGGAAGTATTACCAAG<br><br>L-F1f:<br>AGGAGCCTGAAGCTAAGCTTGTTG | L-R5d :<br>ACCTACCCTCAAGAGCAAAAGTGAGTTCC<br><br>L-R4:<br>CTTCATAAGGTGTGCCAATGTACCTCTTCAT<br><br>L-R4b:<br>GATCAGAGAAACTAAGGTGTCGATAATTC | LongAmp Taq PCR, with annealing/extension time: 4m44s | | | |
| L 301-1738 | L internal | Nucleotide positions 301 – 1738 of L segment (EU044832.1) | 1438 | L-F1:<br>TTTGAGATTGTGCGGCAGCCTGGTGATGG<br><br>L-F1b:<br>GAACTGCTTCTACCACAGCATAGCTGAG<br><br>L-F1c:<br>ACCATGCCCAACAAAACAGCTCACTC | | L1a-R5:<br>GTCTCTTGATCTCGATCAGAAGCTG<br><br>L1a-R6:<br>TCTACTTCAGAATCTTCTTGAAGAGCATA | RT-PCR:<br>SuperScript III One-Step RT-PCR System with Platinum Taq DNA Polymerase, with extension time: 1m50s | L-F1d:<br>ATATCACAACATCAAACATCTGACCGAG<br><br>L-F1e:<br>TTGGCAGCAAGGAAGTATTACCAAG<br><br>L-F1f:<br>AGGAGCCTGAAGCTAAGCTTGTTG | L1a-R3:<br>CTCAAGTAAAGGTTATGTGCATAGGA<br><br>L1a-R4:<br>TGAAATATCCTCAGAGGTCATATTTA | LongAmp Taq PCR, with annealing/extension time: 2m32s | | | |
| L 1-4020*<br>(RT-PCR amplified a sheep sequence non-specifically, but no CCHFV sequence) | L upstream end | Nucleotide positions 1 – 4020 of L segment (EU044832.1) | 4020 | CCHF-L1F:<br>TCTCAAAGATATCAATCCCCCC<br>(10 µM RT primer stock used) | fL0:<br>TCTCAAAGATATCAATCCCCCCTTTACCCA<br><br>fL1:<br>TCTCAAAGATATCAATCCCCCCTTTACCCACGTTG<br><br>fL2:<br>TCTCAAAGATATCAATCCCCCCTTTACCCACGTTGACACAGT<br><br>fL3:<br>TCTCAAAGATATCAATCCCCCCTTTACCCACGTTGACACAGTCAGCTCC | L-R6:<br>TTTATGAGGAAGCCATAAACCAGATTTAAGCCTGA<br><br>L-R7:<br>TACTAAGCCCTGCAAGCATGCCAAAACGAAGCAT | RT:<br>Maxima H Minus First Strand cDNA Synthesis Kit, with reaction temperature: 65 °C<br><br>PCR:<br>LongAmp Taq PCR, with annealing/extension time: 5m | fL4:<br>TCTCAAAGATATCAATCCCCCCTTTACCCACGTTGACACAGTCAGCTCCAGTGGTG | L-R5:<br>ACTGTTTATAACTTCTAGACACCTATACTGCTGCA | LongAmp Taq PCR, with annealing/extension time: 5m | fL5:<br>TCTCAAAGATATCAATCCCCCCTTTACCCACGTTGACACAGTCAGCTCCAGTGGTGGATTTTTGTCTCTGTGGTACCATGGA | L-R4:<br>CTTCATAAGGTGTGCCAATGTACCTCTTCAT | LongAmp Taq PCR, with annealing/extension time: 5m |

* L 1-4020 RT-PCR amplified the expected CCHFV amplicon from cell culture but a sheep-like sequence from tissue homogenates.

**Appendix Table 2. RT-PCR conditions and primer sequences for targeting the strand polarity of CCHFV RNA transcripts**

| Genomic segment from which RNA transcript is derived | Strand polarity of RNA transcript targeted | Genomic location of final amplicon | Size of final amplicon (bp) | Reverse transcription (RT) | | Nested PCR round 1 | | | Nested PCR round 2 | | | Nested PCR round 3 | | |
|---|---|---|---|---|---|---|---|---|---|---|---|---|---|---|
| | | | | RT primer or primer pool (Primer name: Primer sequence) | Reaction conditions | Forward primer (Primer name: Primer sequence) | Reverse primer (Primer name: Primer sequence) | Reaction conditions | Forward primer (Primer name: Primer sequence) | Reverse primer (Primer name: Primer sequence) | Reaction conditions | Forward primer (Primer name: Primer sequence) | Reverse primer (Primer name: Primer sequence) | Reaction conditions |
| S | + | Nucleotide positions 290 – 549 of S segment (DQ133507.1) | 260 | CCHF-SR2: CADGGTGGRTTGAARGC | Maxima H Minus First Strand cDNA Synthesis Kit, with reaction temperature: 50 °C | Nested1-F2-Hoti: TGGACACTTTCACAAACTC | Nested1-R3-Hoti: GACAAACTCCCTGCACCA | LongAmp Taq PCR, with annealing/extension time: 2m24s | Nested2-F3-Hoti: GAATGTGCTTGGGTCAGCTC | Nested2-R2-Hoti: GACATCACAATTTCGCCAGG | LongAmp Taq PCR, with annealing at 60 °C for 30s and extension at 65 °C for 15s | | | |
| | - | | | fS2: TCTCAAAGAAACACGTGCCGCTTACG<br>fS3: TCTCAAAGAAACACGTGCCGCTTACGCCCA<br>fS4: TCTCAAAGAAACACGTGCCGCTTACGCCCACAGT<br>fS5: TCTCAAAGAAACACGTGCCGCTTACGCCCACAGTGTTCTC<br>fS6: TCTCAAAGAAACACGTGCCGCTTACGCCCACAGTGTTCTCTTGAGTGTCTG | | | | | | | | | | |
| M | + | Nucleotide positions 4122 – 4381 of M segment (EU037902.1) | 260 | M-Down-R1: AGCATATTCTTCAATCCATAAAA | | M-Down-F1: TGTGGCTGCACATCATCAACCTG | M-Down-R1: AGCATATTCTTCAATCCATAAAA | | M-Down-F2: AACACCTCTTGGATGTCCTGGGA | M-Down-R2: GCAACCTCAACCAGCACAGTGA | LongAmp Taq PCR, with annealing/extension time: 2m24s | | | |
| L | + | Nucleotide positions 6363 – 7499 of L segment (EU044832.1) | 1137 | CCHF-L1R2: TTCCAGAATGTTTCATCATATTGGGA | | CCHF-L2F-Hoti2: GAGCACCCCAGTGTCAGCTTCAC | CCHF-L1R2: TTCCAGAATGTTTCATCATATTGGGA | | CCHF-L2F-Hoti3: TCAGCTCCTTGGTGCTTACACTACAAAGCAC | CCHF-L1R3: GCATGTTCAACATGTACTGTTGT | | CCHF-L2F-Hoti4: AGACACCCAGAGATACTTGACTGTGG | CCHF-L1R4: ACCCTGACCCATATGGTTGTAACTGTT | LongAmp Taq PCR, with annealing/extension time: 2m24s |

**Appendix Table 3. Visualization of CCHFV RNA and protein in tissues**

| Sheep tissue type | Detection method and target | | | |
|---|---|---|---|---|
| | ISH: Negative-sense RNA (M segment) | ISH: Positive-sense RNA (M segment) | IHC: Protein (N protein) | IHC: Protein (N protein) - deeper recuts through block |
| Sheep 1 Tracheobronchial LN | Weak positive | Weak positive | Negative | |
| Sheep 1 Deep Cervical LN | Negative | Weak positive | Negative | All 5 deeper sections negative |
| Sheep 2 Inguinal LN | Weak positive | Negative | Negative | All 5 deeper sections negative |
| Sheep 2 Gastrohepatic LN | Negative | Negative | Negative | |
| Sheep 3 Gastrohepatic LN | Negative | Negative | Negative | |
| Sheep 3 Inguinal LN | Weak positive (stronger than positive-sense RNA) | Weak positive | Negative | |
| Sheep 4 Inguinal LN | Positive (stronger than positive-sense RNA) | Positive | Positive | |
| Sheep 2 Spleen | Weak positive | Negative | Negative | |

For each tissue one section was stained unless otherwise indicated. LN, lymph node. ISH, in situ hybridization. IHC, immunohistochemistry.

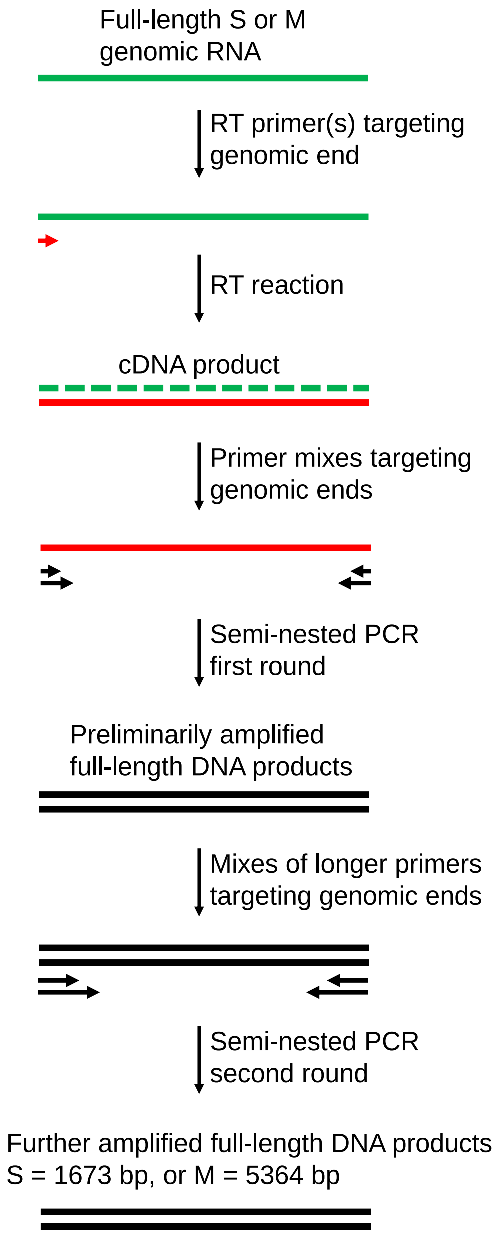

**Appendix Figure 1. RT-nested PCR to amplify full-length CCHFV S or M genomic RNA from sheep tissues.** Diagram is not drawn to scale. Green and red colors specify negative sense and positive sense of CCHFV sequences, respectively. Green dash line and red solid line together represent RNA:DNA hybrid during reverse transcription. Semi-nested PCR rounds used primers targeting the ends of viral genomic segments. Primers of the second round PCR are longer with a different 3' sequence than those of the first round. A new "internal nesting" strategy was applied to each PCR round, in which a mixture of primers with various lengths and 3' sequences were used to target each end of the genomic segment. bp, base pairs. Details of the PCR conditions and primer sequences are provided in Appendix Materials and Methods and Appendix Table 1.

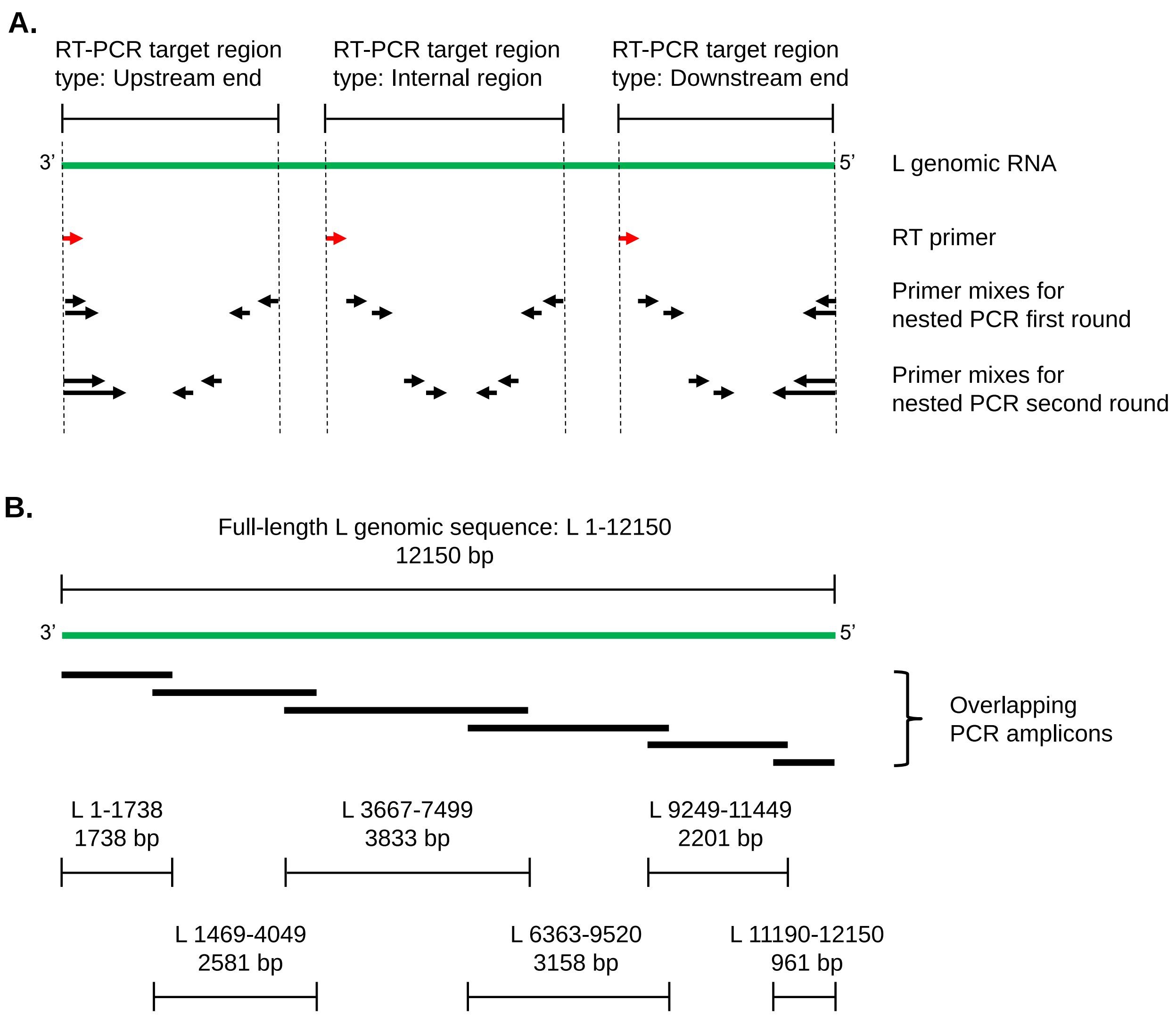

**Appendix Figure 2. RT-nested PCR to amplify CCHFV L genomic RNA sequences from sheep tissues.** The L segment due to its large size was amplified as overlapping amplicons. RT-nested PCR rounds each with internal nesting were performed largely as in Appendix Figure 1 with minor variations: **A.** Diagram illustrates three types of amplicons based on their genomic location. Primers targeting extreme genomic ends were designed similarly to those in Appendix Figure 1 for amplifying full-length S or M segment. Primers targeting genomic internal sites did not share any sequence overlap, as in traditional nested PCR. Some amplicons needed a third PCR round in addition to the two rounds illustrated above. **B.** Genomic locations of the overlapping L amplicons.

```
> Crimean-Congo hemorrhagic fever virus strain Kosovo Hoti segment S, complete sequence, recovered
from tissues of experimentally infected sheep on day 34 post infection
TCTCAAAGAAACACGTGCCGCTTACGCCCACAGTGTTCTCTTGAGTGTCTGCAAAATGGAAAACAAGATCGAGGTGAACAGCAAAGATGAGATGAACAA
ATGGTTTGAGGAGTTTAAAAAGGGAAATGGACTTATGGACACTTTCACAAACTCCTACTCCTTTTGCGAGAATGTACCAAATCTGGATAAGTTTGTGTT
CCAGATGGCCAGCGCCACTGATGATGCACAGAAGGACTCCATCTATGCATCGGCTCTGGTGGAAGCAACCAAGTTCTGTGCACCCATATATGAATGTGC
TTGGGTCAGCTCTACTGGCATTGTGAAGAAGGGGCTTGAGTGRTTTGAGAAGAATTCAGGAACCATCAAATCTTGGGATGAGAACTATGCTGAGCTGAA
GGTTGATGTTCCCAAAATAGAACAACTTGCCAATTACCAACAGGCTGCTCTCAAGTGGAGGAAGGACATAGGTTTCCGTGTCAATGCAAACACGGCAGC
CTTAAGCAACAAAGTCCTTGCAGAATATAAAGTCCCTGGCGAAATTGTGATGTCTGTTAAAGAAATGCTGTCAGACATGATTAGAAGGAGGAATCTAAT
TCTCAACAGGGGCGGTGATGAAAATCCACGCGGCCCAGTGAGCCGTGAACATGTGGAGTGGTGCAGGGAGTTTGTCAAAGGCAAGTACATCATGGCCTT
CAATCCACCTTGGGGGGACATCAACAAATCAGGCCGTTCAGGAATAGCACTTGTTGCAACAGGCCTTGCCAAGCTTGCAGAGACCGAGGGGAAAGGAGT
CTTTGACGAAGCAAAGAAGACCGTGGAGGCTCTCAATGGGTATTTGGACAAGCACAGGGACGAAGTTGACAAAGCAAGTGCCGACAGCATGATAACAAA
CCTCCTAAAGCACATTGCCAAAGCACAAGAGCTTTATAAAAATTCATCTGCTCTTCGTGCACAAGGTGCACAGATTGACACTCCTTTCAGCTCGTTTTA
CTGGCTCTACAAGGCCGGTGTGACTCCAGAGACCTTCCCAACTGTCTCACAGTTCCTCTTCGAACTGGGGAAGCAGCCAAGGGGGACCAAAAAAATGAA
AAAGGCACTCCTGAGCACTCCAATGAAGTGGGGGAAGAAGCTTTATGAGCTCTTTGCTGATGACTCTTTCCAGCAGAACAGAATCTACATGCACCCTGC
TGTGTTGACAGCCGGTAGAATCAGTGAAATGGGTGTCTGCTTTGGAACAATCCCTGTTGCCAATCCTGATGACGCTGCTCAGGGATCTGGACATACCAA
GTCCATTCTCAACCTTCGGACAAGCACAGAGACCAACAACCCATGTGCCAAGACAATTGTCAAATTATTTGAAATCCAAAAAACAGGATTTAACATACA
GGACATGGACATTGTAGCCTCTGAGCACCTGCTGCACCAATCCCTTGTTGGCAAGCAGTCTCCATTCCAAAATGCCTACAATGTCAAGGGCAATGCCAC
TAGTGCCAACATCATCTAGAACTCAAGGCGTTTCACATTCAGCTTTTCTCCTCCTGCATCACTACTTACAGTTATGACTATTAATCACGTTTATTTTAA
TTGCTTATATAATGCTGTTTTGCTAATTTTATCTTGCTATCTTTCATTTCAAATACTTAAAGGGCTGTGCGGCAACGATATCTTTGAGA
```

**Appendix Figure 3. Sequence of CCHFV S segment from sheep tissues.** Full-length S segment genomic RNA was amplified by RT-nested PCR and sequenced by NGS. Sequences from lymph nodes of different sheep were aligned by Clustal W method using DNASTAR Lasergene software. Consensus sequence is shown in positive sense, with standard codes by International Union of Pure and Applied Chemistry (IUPAC). Shaded is nucleotide change in tissues relative to the reference sequence (GenBank accession number DQ133507.1), which was not found in the stock virus used to inoculate animals.

> Crimean-Congo hemorrhagic fever virus strain Kosovo Hoti segment M, complete sequence, recovered from tissues of experimentally infected sheep on day 34 post infection

TCTCAAAGAAATACTTGCGGCACGTCAGTACGTAAGTGTCAACTTTGAGAGAGTGGATTGTGCATCCCGAATGCACCATGCCCATCAACATTATGCATA
CACTATTAGTGTGCTTCATTCTTTACCTACAGCTGTTGGGTCTAGGAGGAGCTCATGGACAGTCAAACGCAACTGAACACAATGGAACAAACACCACCA
CAGCACCCGGCACAAGTCAAAGCCCCAAGCCACCCGCGAGCACAACCCCATCACACGCACCGGAACCATCCACCATCAAACTCACCACACCAACAAGCG
AAACAGAAGGCTCAGGGGAAACGACCCCACCCACAAACACCACCCAGGACTCGCCCCCGCTAGAGACCACACCAGAACGCTCTGCAACAACCGCCACAA
GCACGTCAGGCACAGACAACATGAATTCCACCACACAGATGACCGATAACACCCCCACATCAACAGTCAGCACAAGTCTATCCAGTAGCCCCAGCACGC
CATCCACATCACAAGGCATGCACCATCCCGTAAGGAGTCTATTGTCAGTCTCGAGCCCCAAGACAGCAACAACACCAACACCAATGAGCCCTGGAGAGA
TGAGCTCGGAAACCAGCAGTCAGCACTCAGCCATGAGCAGAACACCAACCCCCCACACAGCAACCCAGGTCTCCACTGAGAACGCCAACCGCAGCACAT
CCAGACAGTCTGAGTCCTCAGCACAGCCGGCAACCCAAAGCCCAGTGACTTCCCCAGCGCAATCAATCCTTCTGATGAGTGCTACTCCCACAACCATTC
AGAACATACATCCCAGCCCAA**T**AAACAGGTCTAAGAGGAACCTTGAGATGGAAATAATTTTAACTCTGTCTCAAGGTCTGAAGAAATACTATGGTAAAA
TACTGAAGCTTCTGCACCTTACCTTGGAAGAGGACACTGAAGGCTTGTTGGAATGGTGTAAGAGAAACCTTGGGTCGAACTGTGATGATGACTTCTTTC
AAAAGAGGATAGAAGAATTCTTCATAACTGGTGAGGGCTACTTCAACGAAGTTCTACAGTTTAAAACACTAAGCACACCAAGCTCCACGGAGCCGTCTC
ATGCCAGGCTACCAACAGCGGAACCCTTCAAATCTTACTTCGCTAAGGGCTTCCTTTCAATAGATTCGGGTTACTTCTCTGCCAAATGTTATCCAAGAT
CATCCACTTCAGGGCTTCAGCTGATCAATGTCACCCAACACCCAGCTAGGATAGCAGAGACACCTGGACCCAAAACAACGAGTCTGAAAACCATCAACT
GCATCAATCTAAGAGCATCTGTCTTCAAAGAACACAGGGAAGTTGAGATCAATGTGCTTCTTCCTCAAATTGCAGTCAACCTCTCAAACTGCCATGTTG
TGATCAATTCACATGTCTGTGATTACTCTTTAGACACTGATGGGCCAGTGAGGCTTCCTCGCATTTACCATGAAGGGACTTTCATGCCAGGAACTTATA
AAATAGTGATAGACAGAAAAAATAAGCTAAATGACCGATGTACGCTAGTCACCAACTGTGTGATAAAAGGAAGAGAAGTTCGTAAGGGCCAATCAGTGC
TGAGGCAGTACAAAACAGAAATTAAGATAGGCAAGGCACCAACAGGCTCTAGGAAACTACTGTCCGAAGAGCCAGGTGATGATTGCATATCAAGGACTC
AGCTATTGAGGACAGAAACAGCAGAGATCCATGACGATAACTATGGTGGTCCAGGTGACAAAATAACTATCTGCAATGGTTCAACCATTGTAGATCAGA
GACTGGGCAGTGAACTAGGGTGTTACACCATCAACAGGGTGAAGTCATTCAAGCTATGCGAAAACAGCGCCACAGGGAAAACCTGTGAGATAGACAGCA
CTCCAGTTAAGTGCAGGCAAGGTTTTTGTTTAAAAATCACCCAAAAGGGAAGGGGCCACGTAAAATTATCTAGAGGCTCAGAGGTTGTCTTGGACGCTT
GCGACTCAAACTGTGAAGTAATGATACCYAAAGGCACTGGAGATATCCTAGTGGACTGTTCAGGTGGACAGCAGCATTTCTTAAAAGACAACTTGATTG
ACCTGGGGTGCCCTCATATCCCACTATTGGGTAAGATGGCCATTTACATTTGCAGAATGTCAAATCATCCTAAAACAACCATGGCTTTTCTCTTCTGGT
TCAGCTTTGGTTATGTGATCACCTGTATATTTTGCAAGGCTCTTTTTTATTCATTGATAATTATTGGGACACTAGGAAAAAAGATCAAGCAATATAGGG
AACTGAAACCTCAAACTTGCACCATATGTGAAACAGCTCCTGTCAATGCAATAGATGCTGAAATGCATGACCTCAACTGTAGTTACAATATATGCCCTT
ATTGTGCATCAAGATTGACCTCCGATGGGCTAGCTAGGCATGTAAAGCAATGCCCTAAGCGAAAGGAGAAAGTCGAGGAGACTGAGCTGTACCTGAACC
TGGAAAGAATTCCTTGGGTCGTCAGAAAGCTATTGCAAGTGTCAGAGTCCACTGGTGTGGCATTGAAACGAAGCAGTTGGCTGATTGTGCTGCTTGTGT
TACTCACAGTTTCATTGTCACCGGTTCAATCAGCACCTGTTGGTCATGGTAAGACGATCGAAACATATCAGACCAGAGAAGGATTCACAAGTATTTGTC
TCTTTATGTTAGGAAGCATCCTATTCATAGTC**C**CTTGCTTGGTGAAAGGGCTGGTTGACAGTGTCAGTGACAGCTTCTTCCCCGGCCTGTCTGTCTGCA
AGACATGCTCCATTGGCAGTATAAATGGCTTTGAAATTGAATCGCATAAATGCTACTGTAGCTTATTTTGCTGCCCCTATTGTAGACACTGCTCTGCTG
ACAGAGAAATTCACCAATTGCACTTGAATATCTGCAAAAAAAGAAAAACGGGGAGCAATGTCATGCTAGCTGTCTGCAAGCGCATGTGCTTTAGGGCAA
CCATAGAAGCAAGCAGAAGAGCCCTGCTCATCCGAAGTATTATCAATACCACTTTTGTAATATGCATTCTAACATTAACAATCTGTGTTGTTAGTACCT
CTGCAGTAGAGATGGAAAACCTTCCAGCAGGCACCTGGGAGAGAGAGGAAGACCTGACAAATTTTTGTCATCAGGAATGCCAGGTAACCGAGACGGAAT
GCCTTTGTCCATACGAAGCTCTTGTGCTCAGGAAACCTCTTTTCTTAGACAGTATAGTCAAAGGTATGAAAAATTTGCTAAACTCAACAAGCTTAGAAA
CAAGCTTATCAATTGAG**A**CACCATGGGGAGCAATAAATGTCCAGTCAACCTTCAAACCAACAGTGTCGACTGCGAACATAGCACTCAGTTGGAGCTCAG
TGGAACACAGAGGCAACAAGATCTTAGTCACAGGCAGGTCTGAGTCAATTATGAAACTAGAGGAAAGGACAGGAATCAGCTGGGATCTAGGCGTAGAAG
ATGCCTCTGAATCCAAATTACTTACCGTCTCTATCATGGATTTGTCACAGATGTATTCCCCAGTTTTTGAGTACTTGTCAGGGGACAGACAGGTTGAAG
AATGGCCCAAAGCAACTTGCACAGGTGACTGCCCACAAAGGTGTGGCTGCACATCATCAACCTGCTTGCATAAAGAATGGCCTCATTCAAGAAACTGGA
GATGTAATCCCACTTGGTGCTGGGGAGTGGGAACTGGCTGCACCTGCTGCGGGGTGGATGTAAAAGACCTTTTTACAGATCACATGTTTGTCAAATGGA
AGGTTGAATATATAAAAACTGAAGCCATAGTGTGTGTTGAGCTCACCAGCCAAGAAAGACAGTGCAGTCTGATTGAGGCAGGTACTAGATTCAACTTGG
GCCCTGTGACTATTACCCTGTCTGAGCCAAGAAACATTCAGCAGAAGCTCCCCCCTGAGATCATTACGTTGCATCCCAAAATTGAGGAAGGCTTCTTTG
ACTTGATGCATGTACAAAAAGTGTTGTCCGCAAGTACAGTGTGTAAACTGCAGAGTTGCACACATGGTATACCAGGAGATCTACAAGTTTACCACATTG
GCAATTTGTTGAAAGGGGATAGAGTGAACGGGCATCTAATTCACAAAATTGAACCACATTTCAACACCTCTTGGATGTCCTGGGATGGTTGTGACTTAG
ACTACTACTGCAACATGGGGGACTGGCCTTCTTGTACATACACAGGAGTGACCCAGCATAATCATGCTGCATTTGTAAACTTGCTCAACATTGAAACTG
ATTACACAAAAACCTTCCACTTCCACTCTAAAAGAGTCACAGCACATGGTGACACACCACAATTGGACTTAAAAGCAAGGCCGACCTACGGTGCAGGAG
AGATCACTGTGCTGGTTGAGGTTGCTGACATGGAGTTGCATACCAAGAAAGTTGAGATATCGGGCTTGAAATTTGCAAGTCTGGCTTGCACAGGTTGCT
ATGCCTGTAGCTCTGGTATCTCCTGTAAAGTCAGGATTCATGTAGATGAACCAGATGAGCTCACAGTGCATGTAAAGAGCAGTGATCCAGATGTGGTTG
CAGCTGGCACAAGCCTCATGGCAAGGAAGCTTGAATTTGGGACAGACAGCACATTCAAGGCCTTTTCAGCTATGCCCAAAACCTCTTTATGTTTTTACA
TTGTGGAGAAAGAATACTGCAAGAGCTGCAGTGAAGATGATACAAAAAAATGTGTTGACACAAAACTTGAGCAGCCACAGAGCATACTAATTGAGCACA
GAGGGACGATAATTGGAAAGCAGAATGATACTTGCACAGCCAAGGCAAGCTGTTGGCTAGAGTCAGTCAAGAGTTTTTTTATGGATTGAAGAATATGC
TTGGCAGTGTTTTTGGTAATTTTTTCATAGGTATTTTGTTATTCCTTGCCCCTTTTGTCCTGTTGGTGCTTTTCTTTATGTTTGGATGGAAAATCTTGT
TTTGCTTTAAATGTTGCAGGAGGACCAGAGGCTTGTTTAAGTATAGACACCTCAAAGACGATGAGGAAACAGGCTACAGAAGGATTATTGAAAGGCTGA
ACAACAAAAAAGGGAAAAATAGACTGCTTGATGGTGAAAGGTTGGCTGATAGGAAAATTGCCGAGCTGTTTTCCACAAAAACACACATTGGCTAACTCA
ACAGAAAGAACCTCGAAGGTTGACAGCAGTATGCCGTCTAGTGCCCTTGCTTATTCTAGTTTCGTTCATCGCATCCATACTTTACAGACACTGCAACAC
ACTGCATCTAGCCTGCCTCATTATATCCACAAATAGGCTCTGTAATGCTTGAAACTGCCCTAGCTCTGCTTGCTCTGACCTTACCTTTGACTGCGTGCC
GCCACTATATCTTTGAGA

**Appendix Figure 4. Sequence of CCHFV M segment from sheep tissues.** Shaded is nucleotide change relative to the reference sequence (GenBank accession number EU037902.1) that was not found in the stock virus. Bolded and underlined are nucleotide changes that were carried over from the stock virus.

> Crimean-Congo hemorrhagic fever virus strain Kosovo Hoti segment L, complete sequence, recovered from tissues of experimentally infected sheep on day 34 post infection

TCTCAAAGATATCAATCCCCCCTTTACCCACGTTGACACAGTCAGCTCCAGTGGTGGATTTTTGTCTCTGTGGTACCATGGACTTCTTGAGGAACCTTGATTGGACTCAGGTGATTGCTGGTCAGTATGTGACCAACCCTAGGT
TCAATATCTCCGATTACTTTGAGATTGTGCGGCAGCCTGGTGATGGGAACTGCTTCTACCACAGCATAGCTGAGTTAACCATGCCCAACAAAACAGCTCACTCATATCACAACATCAAACATCTGACCGAGTTGGCAGCAAGGA
AGTATTACCAAGAGGAGCCTGAAGCTAAGCTTGTTGGCCTGAGCCTGGAAGATTACCTTAAGAGGATGCTGTCTGACAACGAGTGGGGTTCAACTCTTGAGGCATCCATGCTAGCCAAAGAAATGGGCATTACAATAATCATTT
GGACTGTTGCTGCCAGTGATGAAGTGGAGGCAGGCATAAAGTTCGGAGACGGTGATGTGTTTACAGCTGTTAACCTTCTGCACTCTGGGCAAACACATTTTGATGCTCTCAGAATACTGCCACAGTTTGAAACAGACACAAGGG
AAGCCTTGAGTTTGGTGGATAAGGTTATGGCTGTGGATCAGTTGACATCATCCTCTAGTGATGAGCTGCAAGACTATGAGGATCTTGCTTTAGCACTCACAAGTGTGGAAGAACCGTATAGGCGGTCAAGCTTGGATGAAGTCA
CTCTGTCCAGGAAACAAGCAGAGATATTGAGGCAGAAGGCATCTCAGTTGTCTAAATTGGTCAATAAAAGTCAGAACATACCGACTAGAGTCGGCAGGGTCTTGGACTGCATGTTTAACTGTAAACTATGTGTTGAGATATCAG
CTGACACTTTAATTCTAAGGCCAGAATCCAAAGAGAGAATTGGCGAAGTCATGTCATTGCGGCAGCTGGGGCACAAATTGCTGACACGAGACAAACAAATTAAGCAAGAATTCTCTAGGATGAAACTTTACGTCACCAAAGATT
TGCTAGACCATCTAGATGTTGGTGGGCTGTTGAGGGCTGCTTTCCCTGGGACAGGAATAGAAAGGCATATGCAGCTGCTACACTCCGAGATGATACTGGACATCTGTACTGTATCACTTGGCATCATGTTGTCAACATTCCTAT
ACGGCTCTAACAATAAGAACAAGAAGAAGTTCATTACCAACTGTCTGCTCAGTACAGCTTTGTCCGGAAAAAAGGTGTATAAGGTTCTYGGCAACCTAGGAAATGAACTGTTGTACAAGGCACCCAGAAAGGCCTTAGCAACTG
TCTGTAGTGCCTTGTTTGGGAAGCAGATAAACAAGCTTCAGAACTGCTTCAGAACCATAAGCCCCGTCAGCTTACTTGCATTAAGAAATCTGGACTTTGACTGTCTAAGTGTGCAAGACTATAATAACATGATAGAAAGTATGT
CCAAATTGGATAACACTGATGTTGAATTCAACCACAGGGAGATAGCTGATCTCAATCAATTAACTTCTCGGCTCATTACATTGAGAAAAGAGAAAGACACTGACCTCCTCAAACAATGGTTTCCTGGAAGTGACCTCACCCGCA
GAAGTACCAGGAATATCGCAAACGCAGAGGAATTTGTCATTTCTGAGTTCTTCAAAAGGAAGGACATTATGAAATTCATCAGCACCTCAGGAAGGGCAATGAGTGCAGGCAAGATTGGAAATGTCCTTTCCTATGCACATAACC
TTTACTTGAGTAAGTCTAGTTTAAATATGACCTCTGAGGATATTTCACAGCTTCTGATCGAGATCAAGAGACTGTATGCTCTTCAAGAAGATTCTGAAGTAGAGCCGATAGCCATAATTTGTGATGGCATAGAAAGCAATATGA
AACAGTTGTTTGCTATACTGCCTCCTGACTGTGCAAGAGAGTGTGAGGTCCTCTTTGATGACATAAGAAACTCTCCAACGCACAGTACAGCTTGGAAGCATGCCCTTCGGTTAAAAGGGACTGCATACGAAGGTCTGTTTGCAA
ACTGCTACGGATGGCAATACATCCCAGAAGATATCAAACCAAGCCTGACCATGTTGATACAGACTTTGTTTCCTGACAAGTTTGAAGATTTCTTGGATCGAACCCAGTTGCATCCAGAGTTCAGAGATTTAACTCCTGACTTTT
CGCTTACACAGAAGGTTCACTTCAAAAGAAATCAGATACCCAGTGTTGAAAATGTGCAAATCTCCATTGATGTAACACTGCCTGAGTCTGTGGAGGCAGTGCCGGTGACAGAAAGAAAAATGTTCCCTCTGCCTGAAACTCCGC
TGAGTGAGGTGCATTCAATAGAGCGCATTATGGAAAATTTTTCCCGCCTTATGCATGGAGGAAGACTTTCAGCCAAGGAAAAAGATGGAGACCTTGTAGAACAAGGCAGCCAGCAAAGCGCCGTTGAACACGAGAGTCCAAGCA
TCTCGGCCTTCAAGGACTATGGAGAAAGAGGAATAGTTGAGGAGAATCACATGAGGCTTAGTGAAGAAGATCAACTAGAAACAAGGCAGCTGCTGTTGGTGGAAGTTGGTTTTCAAACTGATATCGATGGGAAAATAAGGACAG
ACCACAAGAAATGGAAGGATATTTTAAAGCTGTTAGAGTTGCTAGGAATCAAATGCTCTTTTATCGCCTGTGCCGACTGTTCGTCTACACCGCCAGATAGGTGGTGGATTACGGAGGACAGAGTACGAGTCTTAAAGAATTCAG
TCAGTTTCCTCTTTAATAAACTCTCCAGGAACTCACCCACAGAAGTAACTGACATAGTTGTTGGAGCTATAAGTACTCAAAAGGTTAGGAGTTACCTAAAGGCAGGAACTGCAACAAAAACCCCTGTGTCGACAAAAGACGTTC
TGGAGACTTGGGAAAAAATGAAAGAGCACATACTAAATAGGCCAACAGGATTAACACTGCCTGCCAGTTTGGAACAGGCTATGCGCAAAGGACTAGTTGAAGGTGTGGTCATCTCTAAGGAAGGTTCTGAATCATGCATCAACA
TGTTGAAAGAAAATTTGGACCGAATAACTGATGAATTCGAGCGAACAAAATTTAAACATGAACTTACTCAGAATGTTACCACTAGTGAAAAGATGCTATTTAGCTGGTTGAGTGAAGATATCAAATCATCGAGATGTACTGAGT
GCCTCACCAACATAAAAAAAACTGTTGACGAGACTGCCAACCTGTCTGAGAAGATTGAGCTGCTTGCATATAATCTTCAACTCGTTAGTCATTGCGGCAACTGTCACCCCAATGGTGTGAGCATTAGCAATACATCTAATGTAT
GCAAGAGGTGCCCTAAAATAGAAGTGGTTAATCATTGTGAAAATAAAGGCTTTGAGGACAGCAATGAATGCTTAACAGATCTAGACAGGCTTGTTAGGCTCACATTACCAGGAAAAACTGAGAAGGAGAGAAGAGTCAAACGCA
ATGTGGAAT**R**CCTTATAAAACTGATGATGAGTATGTCAGGCATTGATTGCATAAAATATCCCACAGGACAGCTTATCACCCATGGAAGAGTGAGTGCAAAACATAACGATGGAAATTTGAAAGATAGGAGTGAGGACGATCAAA
GACTAGCCGAGAAGATAGACACTGTTAGAAAAGAGCTTTCTGAATCCAAACTAAAAGATTATTCAACTTATGCAAGAGGGGTGATTTCAAATTCACTGAAAAACCTCTCAAAGCAAGGCARGTCAAAGTGTTCTGTGCCGAGAT
CTTGGCTTGAAAAGATATTGTTTGACTTAAAAGTGCCCACTAAAGATGAAGAAGTGCTTATAAACATTAGAAATTCATTGAAAGCTAGATCTGAGTTTGTTAGAAACAACGATAAACTACTCATAAGGTCAAAAGAAGAATTAA
AAAAATGTTTCGATGTGCAATCATCTAAACTAATGAAAAACAAGCAACCTGTGCCTTTTCAAGTTGACTGCATACTGTTTAAAGAAGTGGCAGCTGAGTGCATGAAGAGGTACATTGGCACACCTTATGAAGGAATTATCGACA
CCTTAGTYTCTCTGATCAATGTTCTGACAAGGTTCACCTGGTTTCAAGAGGTGGTACTATATGGTAAGATATGTGAAACCTTTCTAAGATGTTGTACAGAATTTAATAGGTCAGGGGTTAAACTAGTCAAGGTAAGGCACTGTG
ACATAAACCTATCAGTCAAACTACCATCGAACAAAAAGAGAACATGTTATGTTGTATATACAGTGGCAACATGGAGCTCTTACAAGGACCTTTCTATTTGAACAGGAGACAAGCTGTCCTTGGATCTTCATACCTTTATATTG
TCATTACACTTTACATACAAGTATTGCAGCAGTATAGGTGTCTAGAAGTTATAAACAGTGTAAATGAAAAGACATTGCAAGACATTGAAAATCACTCCATGACTTTGCTAGAAGATGCATTCAAGGAACTCACTTTTGCTCTTG
AGGGTAGGTTTGAAGAATCTTACAAAGTACGGACCTCTAGGTGTAAAGCCAGCGGGAACTTCTTAAACAGGAGTAGTAGAGACCACTTTATAAGCATTGTCTCAGGCTTAAATCTGGTTTATGGCTTCCTCATAAAGGATAACT
TATTGGCTAATTCTCAACAGCAGAATAAACAACTTCAAATGCTTCGTTTTGGCATGCTTGCAGGGCTTAGTAGGCTTGTTTGCCCTAATGAGCTAGGGAAAAAGTTTTCAACAAGTTGTAGAAGGATTGAAGACAACATTGCAA
GACTTTACCTACAAACATCTATATATTGTTCAGTCAGAGACGTGGAGGACAATATTAAACACTGGAAGCAAAGAGATCTATGCCCTGAAGTGACCATTCCGTGCTTTACAGTCTATGGCACCTTTATCAACAGCGACAGACAGC
TTATCTTTGACATTTACAATGTGCACATATACAATAAAGAAATGGATAATTTTGACGAAGGATGTATCAGCGTCTTGGAAGAAACAGCAGAAAGACATATGCTTTGGGAACTTGATCTGATGAACTCACTTTGCTCTGACGAAA
AAAGAGATGCTAGAACCGCAAGATTATTACTAGGTTGCCCAAATGTGAGGAAAGCTGCAAACAAAGAAGGGAAAAAGCTATTAAAGTCAAACAGCGACACATCCACTGATACACAAAGCATTGCTTCTGAGGTGTCAGACAGAA
GGTCTTATA**R**CTCAAGCAAGAGTAGAATCCGTAGTATTTTTGGCAGGTACAACTCTCAGAAAAAACCATTTGAATTAAGGTCAGGCCTTGAAGTCTTTAATGACCCTTTCAATGATTATCAGCAGGCAATAACAGACATTTGTC
AATTTTCTGAGTATACACCAAACAAAGAGAGCATTTTGAAGGACTGTCTTCAAATCATACGGAAGAACCCTAGCCACACAATGGGTTCTTTTGAGCTGATCCAGGCAATCTCAGAATTCGGTATGAGCAAATTTCCTCCAGAGA
ATATAGACAAGGCAAGGAGAGACCCGAAGAACTGGGTCAGCATCTCTGAAGTTACTGAAACGACAAGTATAGTTGCATCACCCAAAACTCACATGATGCTCAAAGATTGCTTTAAAATTATACTAGGTACTGAGAATAAGAAGA
TAGTTAAAATGCTTAGAGGGAAGCTAAAAAAACTTGGTGCTATCTCAACAAACATAGAGATTGGGAAAAGGGATTGCTTGGATCTACTCAGCACGGTAGATGGTCTGACAGACCAACAGAAAGAAAACATTGTAAACGGGATAT
TTGAGCCTTCAAAGCTGTCCTTCTATCACTGGAAAGAGTTGATCAGAAAGAACATTGATGAAGTTTTACTTACTGAAGATGGAAATCTAATCTTCTGCTGGCTAAAAACAATCTCTTCTTCAGTCAAAGGAAGTTTGAAGAAGA
GACTCAAGTTCATGAATGTAAATTCCCCGGAACTGATGCCAGAGAACTGTCTCTTTTCCAGTGAAGAATTCAATGAGTTGATCAAACTGAAGAAGCTCCTCCTCAACGAGCAACAAGATGAGCAGGAGCTGAAACAAGACCTTT
TGATATCCTCTTGGATTAAGTGCATAACAGCCTGTAAAGATTTTGCAAGCATCAATGACAGGGTTCAGAAATTCATTCACCACCTTTCTGAAGAGCTGTATGACATAAGGCTGCAGCACCTGGAACTGTCAAAGCTTAAACAAG
AGCACCCCAGTGTCAGCTTCACTAAGGAAGAGGTCCTAATAAAGCGACTAGAGAAAAATTTCCTAAAGCAGCACAATCTGRAAATTATGGAGACAGTAAACCTTATATTCTTTGCTGCTCTTTCAGCTCCTTGGTGCTTACACT
ACAARGCACTAGAGTCTTATCTAGTAAGACACCCAGAGATACTTGACTGTGGATCTAAAGAGGACTGTAAGCTCACTTTGCTTGATCTGTCAGTTTCSAAGCTCTTAGTCTGTTTGTATCGAAAAGAAGATAATGAACTAACAA
ATAGCTCAAGTTTGAAACTTGGGTTTTTAGTAAAGTATGCAGTTACCCTATTCACATCCAATGGTGAACCTTTTTCGCTTAGTCTGAATGATGGGGGTTTAGATCTTGACCTGCATAAGACTACTGACGAAARGCTACTACATC
AGACAAAGATAGTTTTTGCTAAAATTGGTCTGTCCGGGAATAGTTATGATTTCATCTGGACTACTCAAATGATAGCAAATAGCAACTTTAATGTTTGCAAGAGATTGACAGGAAGGAGTACCGGAGAAAGGCTTCCAAGAAGTG
TCAGAAGCAAGGTCATTTATGAGATGGTAAAGCTAGTAGGTGAGACAGGTATGGCAATACTACAACAATTAGCTTTTGCACAGGCACTAAATTACGAACACCGTTTCTATGCAGTCTTAGCACCTAAGGCACAGTTAGGAGGAG
CAAGAGATCTGTTGGTGCAAGAAACTGGGACTAAAGTCATGCATGCAACAACTGAAATGTTCAGTAGAAACCTTTTAAAGACAACATCAGATGATGGCCTTACCAACCCGCATCTTAAAGAAACAATCCTTAATGTGGGATTAG
ACTGCCTTACCAAYATGCGAAATCTTGATGGTAAACCCATAAGTGAAGGTAGCAACTTGGTCAATTTCTACAAGGTCATATGCATCTCGGGTGATAACACTAAATGGGGCCCGATACACTGCTGTTCCTTCTTTTCCGGCATGA
TGCAGCAGGTTCTAAAAAATGTTCCAGATTGGTGTTCATTTTATAAATTAACATTTATTAAGAACTTGTGTAGGCAGGTRGAGATACCTGCTGGTAGCATTAAGAAGATCTTAAATGTCCTTAGATACAGGTTGTGCAGCAAAG
GAGGTGTAGAGCAATATAGTGAAGAGGATCTAAGGAAACTGTTGGCAGATAATTTAGACAGTTGGGATGGAAATGACACAGTTAAGTTTTTAGTTACAACCTATATAAGCAAAGGACTCATGGCGTTGAACAGTTACAACCATA
TGGGTCAGGGTATTCACCATGCAACATCTTCAGTATTAACTTCTCTAGCTGCTGTGCTCTTTGAGGAGCTGGCAATTTTTTATCTTAAGAAAAGCTTACCCCAGACAACAGTACATGTTGAACATGCGGGCAGTTCAGATGATT
ACGCAAAGTGTATAGTGGTAACTGGTATACTATCCAAAGAGCTCTACTCCCAATATGATGAAACATTCTGGAAACATGCCTGTAGACTTAAGAACTTCACAGCTGCAGTACAAAGATGTTGTCAAATGAAAGATAGTGCCAAGA
CTTTAGTGGGCGACTGTTTCCTTGAATTTTACAGCGAGTTTATGATGGGCTACAGAGTAACCCCTGCTGTGATCAAATTCATGTTCACTGGACTCATAAACAGCTCTGTGACCTCACCTCAGAGTTTGATGCAGGCATGCCAAG
TTTCGTCCCAACAAGCAATGTATAATAGCGTTCCTCTTGTCACCAATACTGCCTTCACACTATTAAGGCAACAAATTTTCTTTAACCATGTTGAGGATTTTATCAGAAGGTACGGCATGTTAACCCTTGGGACTTTGTCTCCCT
TTGGTAGGCTGTTTGTACCGACCTACTCTGGGTTAGTTAGCTCAGCAGTTGCCTTGGAAGATGCTGAAGTCATTGCTAGAGCAGCTCAAACACTTCACATGAACAGTGTATCTATACAGTCAAGCAGCTTGACCACATTGGACA
GCTTAGGTTGCAGTAGGACAAGTTCCACAGTTGAAGATAGCAGCAGTGTGAGCGACACCACCGTTGCTTCTCATGATTCAGGATCGTCATCCTCCAGCTTCTCTTTTGAGCTCAATAGGCCTCTGTCTGAAACTGAACTACAAT
TCATTAAAGCATTAAGTAGCTTCAAATCCACACAAGCCTGTGAAGTAATCCAGAATAGGATCACAGGTCTTTATTGCAATAGCAATGAGGGGCCTCTTGATAGGCACAATGTTATTTATAGCAGCAGAATGGCAGATTCTTGCG
ATTGGCTAAGAGATGGTAAGAGAAGAGGGAATCTGGAACTAGCAAATAGGATTCAGTCTGTGCTGTGTGTTCTGATAGCAGGATACTACAGATCATTTGGAGGTGAAGGAACCGAGAAACAGGTAAAGGCATCATTGAATAGAG
ATGACAACAAAATCATTGAGGATCCCATGATACAATTAATTCCAGAGAAGCTGAGGAGAGAGTTGGAGAGGTTAGGTGTTTCTAGAATGGAAGTTGATGAACTGATGCCAAGCATCAGCCCTGATGACACCTTAGCCCAGCTTG
TGGCAAAAAAGCTTATTAGCCTCAATGTTTCAACGGAAGAGTACTCGGCTGAAGTGTCTAGACTTAAACAAACATTAACAGCCAGAAATGTTCTGCATGGGTTGGCTGGAGGAATTAAGGAGCTCTCGCTCCCAATATACACGA
TATTCATGAAATCTTACTTCTTCAAAGACAACGTCTTCTTGTCGCTAACAGACAGATGGTCTACCAAGCACAGCACAAACTATCGTGATAGTTGCGGCAAACAATTAACCGGTAGAATAATTACCAAGTACACTCACTGGTTGG
ACACCTTTCTAGGTTGCTCGGTCTCCATTAACAGGCATACTACTGTTAAAGAGCCCTCCCTGTTCAATCCAAATATCAGGTGTGTTAACCTAATCACATTTGAAGACGGTCTAAGAGAACTATCAGTAATACAGAGTCATCTCA
AAGTCTTTGAAAATGAATTTACCAACTTAAACCTTCAGTTCTCCGATCCAAACAGACAGAAACTCAGAATAGTTGAGTCTAGACCTGCGGAATCTGAGCTGGAGGCGAACCGTGCAGTAATTGTTAAAACCAAACTGTTCTCGG
CAACTGAACAAGTACGACTATCCAACAATCCTGCAGTTGTCATGGGCTATTTATTGGATGAATCAGCAATTTCTGAAGTCAAACCTACCAAAGTTGACTTCTCAAACTTACTTAAAGATCGCTTTAAAATAATGCAGTTTTTCC
CCTCAGTGTTCACTTTGATCAAGATGTTGACAGATGAGTCATCGGATTCAGAAAGGAATGGCCTTAGTCCAGACTTGCAACAGGTTGCAAGATACTCAAACCATTTAACCCTGCTCAGCAGGATGATACAACAAGCAAAGCCAA
CTGTGACGGTTTTCTACATGCTGAAAGGTAACTTGATGAACACAGAGCCGACAGTTGCTGAGCTTGTCAGCTATGGTATAAAGGAAGGTAGGTTTTATAGGCTCTCAGACACTGGAATCGATGCAAGCACATATTCTGTGAAAT
ACTGGAAAATTCTTCACTGTATTTCTGCCATTGGATGTTTGCCTTTGAGTCAAGCAGACAAGTCTTCACTACTTATGAGTTTCTTAAATTGGAGGGTAAACATGGACATCAGAACTTCTGACTGTCCACTATCTAGTCACGAGG
CAAGTATACTGAGTGAGTTTGATGGACAAGTTATCGCTAACATACTTGCAAGTGAATTGAGCTCTGTAAAACGAGATTCTGAGCGTGAAGGGTTAACTGATCTCCTTGATTATCTGAACTCACCCACTGAACTATTGAAGAAGA
AGCCCTATTTAGGGACAACTTGCAAGTTCAACACCTGGGGAGACTCAAATAGATCTGGGAAATTCACATACAGCAGCAGATCTGGAGAATCCATTGGAATCTTCATTGCAGGAAAATTGCACATCCATCTCTCATCTGAGTCCA
TTGCCTTATTGTGTGAAACTGAAAGACAAGTGCTATCTTGGATGAGCAAAAGGAGAACTGAGGTGATGACCAAAGAACAGCATCAATTGTTCCTGAGCCTCCTCCCTCAGTCACATGAATGTCTACAAAAGCACAAGGACGGCA
GCGCACTTTCAGTCATACCTGACAGCAGCAACCCCCGACTACTCAAATTTGTACCTCTTAAAAAAGGTTTAGCAGTGGTGAAAATCAAAAAACAAATTTTAACAGTGAAGAAACAGGTCGTGTTTGATGCTGAGAGTGAGCCTA
GACTGCAGTGGGGACATGGTTGCTTGTCCATTGTTTACGATGAAACTGACACTCAAACCACATACCACGAGAATCTTTTGAAAGTGAAGCAGCTTGTTGACTGCTCTACAGATAGGAAGAAGCTTTTGCCCCAATCAGTGTTTT
CTGATTCCAAAGTCGTTCTCTCAAGGATCAAATTCAAGACAGAGCTTCTCCTTAATTCACTGACACTGCTCCACTGTTTCTTAAAACATGCTCCTAGTGATGCCATAATGGAGGTAGAGAGTAAGAGCAGCTTGCTGCACAAGT
ACCTCAAATCAGGAGGTGTCAGACAGCGAAACACTGAAGTACTCTTCAGAGAGAAGTTAAACAAGGTTGTCATAAAAGACAATCTCGAACAAGGTGTAGAGGAGGAGATTGAGTTTTGCAACAACCTGACCAAGACTGTTTCAG
AGAACCCGCTACCACTCAGCTGTTGGTCTGAAGTTCAAAACTACATTGAAGATATAGGCTTCAACAATGTGCTTGTTAACATTGACAGAAACACAGTGAAAAGTGAACTTTTGTGGAAATTTACGTTAGACACTAATGTAAGCA
CCACAAGCACCATTAAAGATGTGAGGACACTGGTATCCTACGTCAGTACTGAAACCATCCCCAAGTTCTTGCTTGCATTCCTTCTCTACGAAGAAGTATTGATGAACTTAATCAACCAGTGTAAGGCAGTTAAGGAACTCATCA
ACAGTACAGGACTTTCAGACTTGGAACTAGAAAGCTTGCTTACTTTTTGTACTTTCTATTTCCAAAATGAATGCAGTAGGAGAGATGGACCTAGATGTTCCTTTGCAGCACTATTAAGCCTAGTTCATGAGGATTGGCAAAAGA
TAGGGAAAAATATCCTTGTTCGTGCAAACAATGAGCTAGGCGATGTGTCACTGAGGGTTAATATTGTCTTGGTACCCCTCAAGGACATGTCCAAACCTAAGCCTGAGAGAGTGGTCATAGCTAGGAGGTCACTAAGTCATGCAC
TATCCTTAATGTTTCTGGACGAGATGTCATTGCCTGAGCTAAAGTCTTTGTCCGTGAACTGCAAAATGGGGAACTTTGAGGGGCAGGAGTGCTTTGAGTTTACCATTCTAAAGGACAACAGCACAAGGCTAGACTACAATAAGT
TGATTGACCATTGTGTGGACATGGAGAAAAAAAGGGAAGCAGTTAGAGCAGTAGAAGATTTAATTCTGATGTTAACAGGCAAGGCAGTCAAACCTAGTACCGTGACACAAATTGAACACAAGGACGAGCAGTGTCAAGAGCAAA
TAAGCTTGGATGACCTAATGGCAAGTGATACAGCGATAGACTTGCCTGACAGGGAAGCAGAAGCCCTTAAAACAGGAAACCTTGGCTTTAACTGGGATTCAGATTGAACATGCCATCTGTACGAGCCATTAATGTCTTTCAATG
TCCCAGGGGCAAACGAGACAGTCTATACCATAACAATTGCCAACATCAAAGCTCTCAAACGAGCTAGCTGCTCTGCATTTCTCACAGTCAACTCAGCTGCCCCTATTGATCTATTTTGCCATCCTACCGTTCTTCTTTCTCTAC
TATTCATACCAGTTTCATTATTTAAAGGGTGTGGGGGGAACGATTTCTTTGAGA

**Appendix Figure 5. Sequence of CCHFV L segment from sheep tissues.** Shaded are nucleotide changes relative to the reference sequence (GenBank accession number EU044832.1) that were not found in the stock virus. Bolded and underlined are nucleotide changes that were carried over from the stock virus.

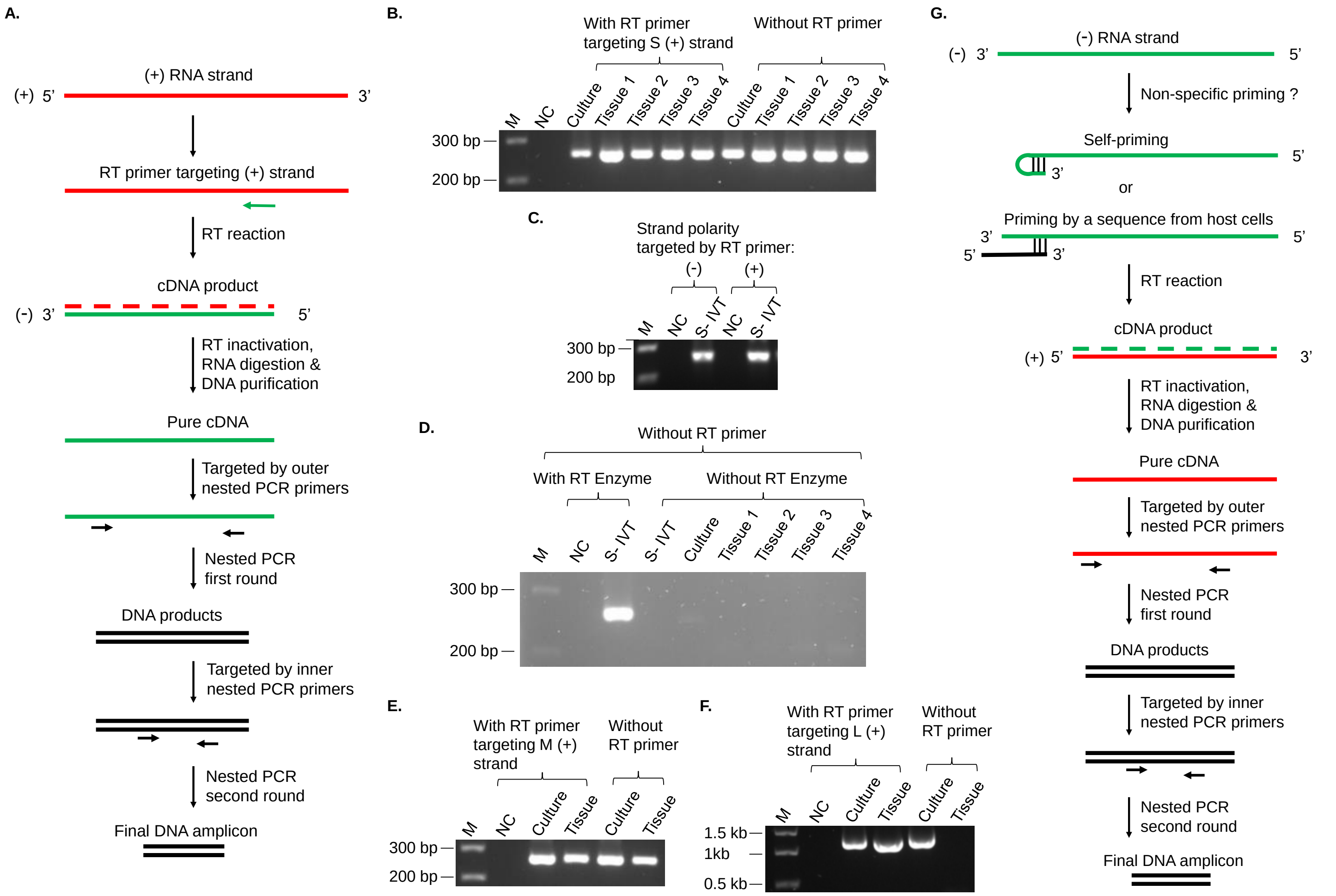

**Appendix Figure 6. RT-nested PCR targeting strand polarity of CCHFV RNA transcripts in total RNA extracts from sheep tissues. A.** Strand-specific RT-nested PCR procedure exemplified for targeting positive-sense transcripts. Red and green colors represent positive sense (+) and negative sense (-) of CCHFV sequences, respectively. The RT reaction was treated with heating to inactivate the RT enzyme and with RNases to remove RNA, including RNase H digestion of the hybrid RNA strand attached to cDNA and RNase A digestion of free RNA. This was followed by a column-based affinity purification of cDNA, which denatures and washes off proteins and enriches DNA. The purified cDNA was then subjected to nested PCRs. Details of the RT and PCR conditions and primer sequences are provided in Appendix Materials and Methods and Appendix Table 2. **B.** Agarose gel of PCR products from RT-nested PCR for detecting S segment transcripts, in the presence or absence of an RT primer aimed at the positive sense. M, marker for DNA molecular sizes, with base pair (bp) numbers labelled on the left. NC, negative control without RNA template. Culture, supernatants of SW-13 cell culture infected with CCHFV. Tissue, lymph node. **C.** RT-nested PCR using an RT primer aimed at an S segment sequence of either negative or positive sense. S-IVT, S segment genomic RNA produced by in vitro transcription, of negative sense only. **D.** RT-nested PCR without RT primer, and with or without RT enzyme, for detecting S segment transcripts. **E and F.** RT-nested PCR for detecting M or L segment transcripts, in the presence or absence of an RT primer aimed at the positive sense. **B – F.** All the PCR products were confirmed by sequencing to be of expected CCHFV sequences. **G.** Hypothetical mechanisms for impaired strand-specific detection. In RT-nested PCR aimed at a positive-sense viral transcript in total RNA extracts, a negative-sense RNA can be non-specifically amplified due to false priming of the RT reaction. The 3' end of the template RNA may fold back to form a primer suitable for the RT. Priming may also be possible by a host cell sequence. Once the negative-sense RNA is converted to cDNA, it is readily amplifiable by a pair of PCR primers irrespective of its polarity.

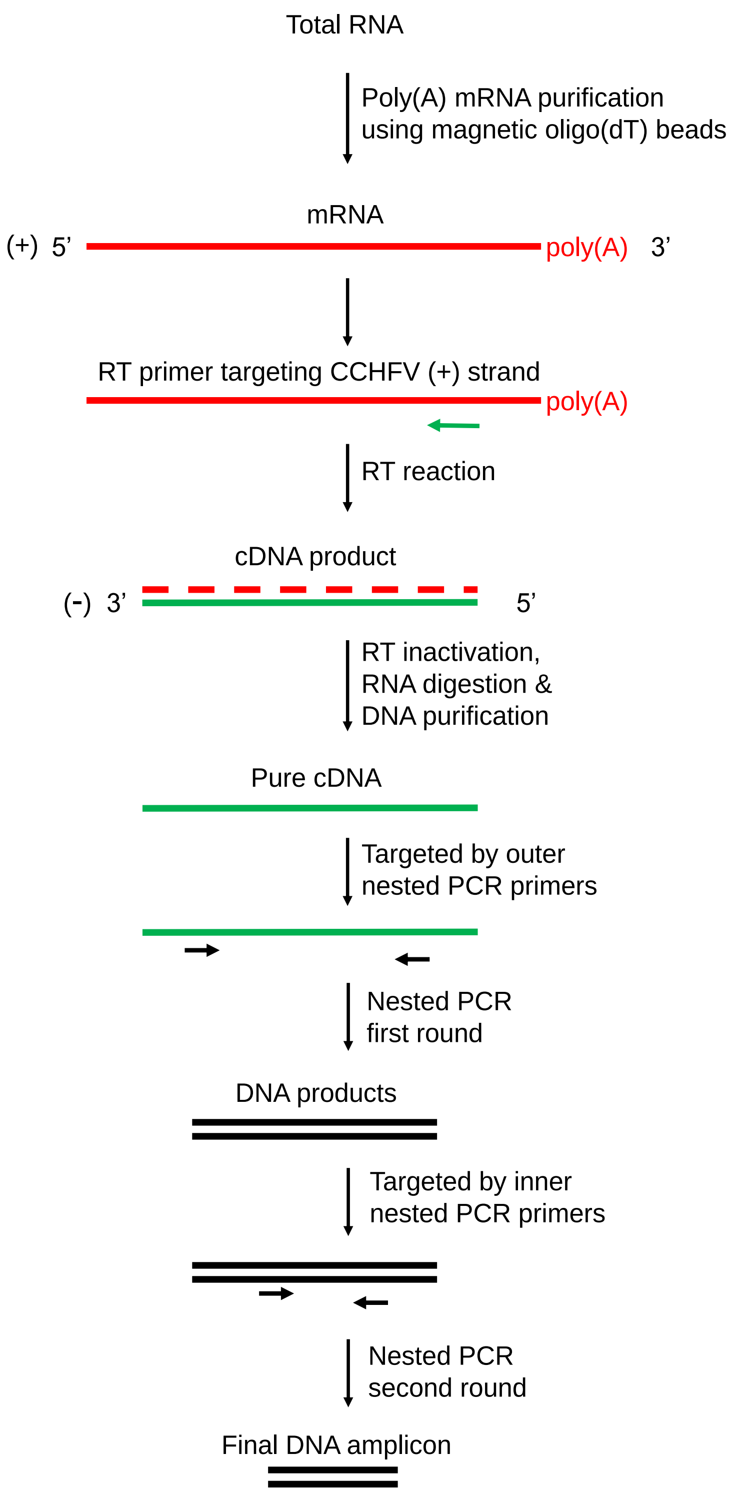

**Appendix Figure 7. RT-nested PCR of CCHFV mRNA transcripts from sheep tissues. A.** mRNA transcripts, with a poly(A) tail, were pulled down using magnetic oligo(dT) beads, while all other components from total tissue RNA extracts including negative-sense, non-poly(A) CCHFV RNAs were washed off. The purified mRNA transcripts were then analyzed by positive sense-targeting RT-nested PCR with RT and PCR conditions and primers detailed in Appendix Table 2.

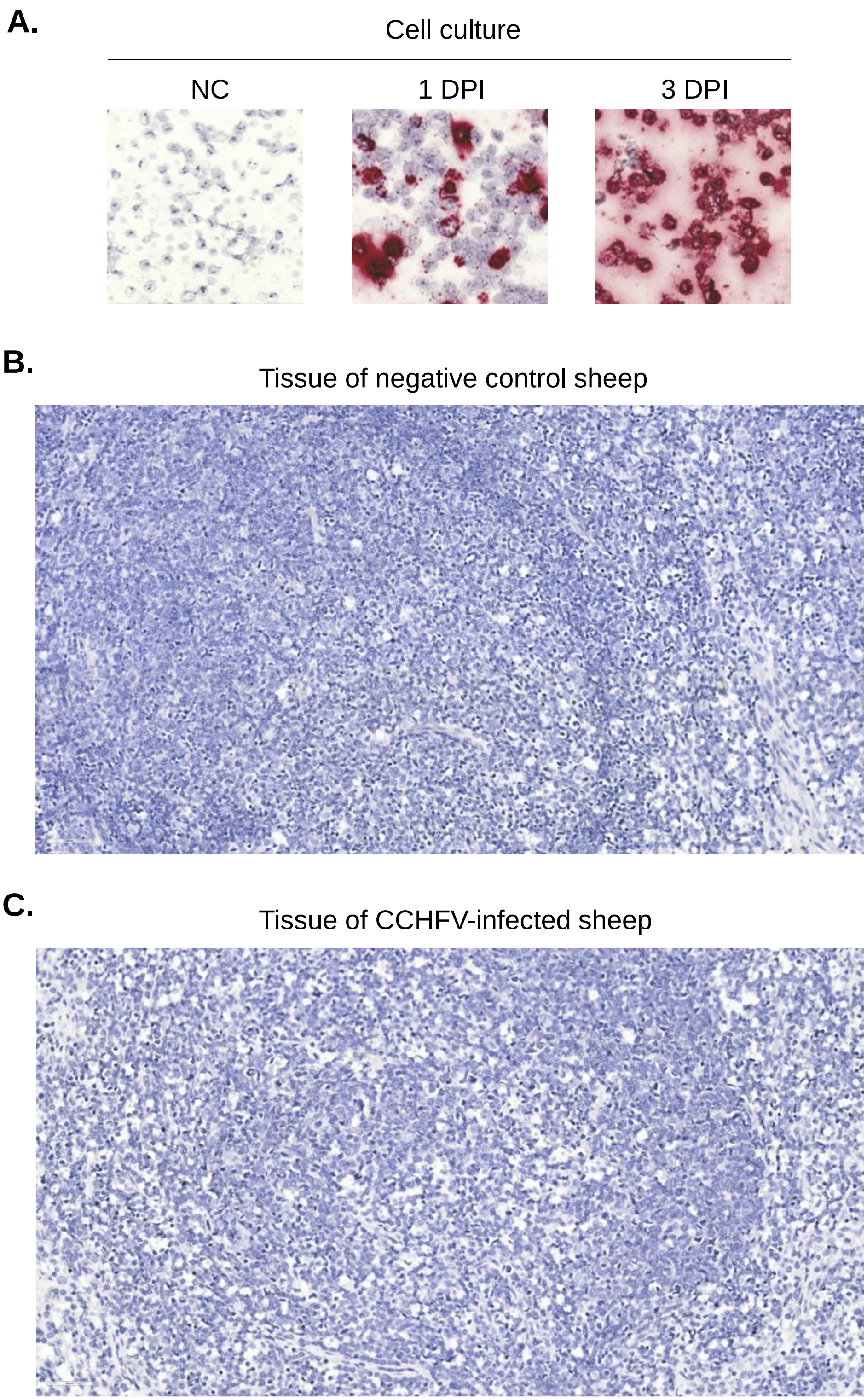

**Appendix Figure 8. Establishment of in situ hybridization (ISH) for negative-sense CCHFV RNA (M segment). A.** Images of control staining using cell pellets from CCHFV cell cultures. NC, negative control cell culture without CCHFV. 1 DPI and 3 DPI, positive control cell cultures with CCHFV on 1 and 3 day(s) post-infection, respectively. **B.** Image of CCHFV-negative control staining using a lymph node tissue from a sheep infected with foot-and-mouth disease virus. **C.** Image representing negative staining in tissues from CCHFV-infected sheep. Image representing positive staining is shown in Figure 2A. A summary of results from multiple sheep tissues are provided in Appendix Table 3. Images were taken at 40× magnification and scaled down in this figure to 12% relative to original size.

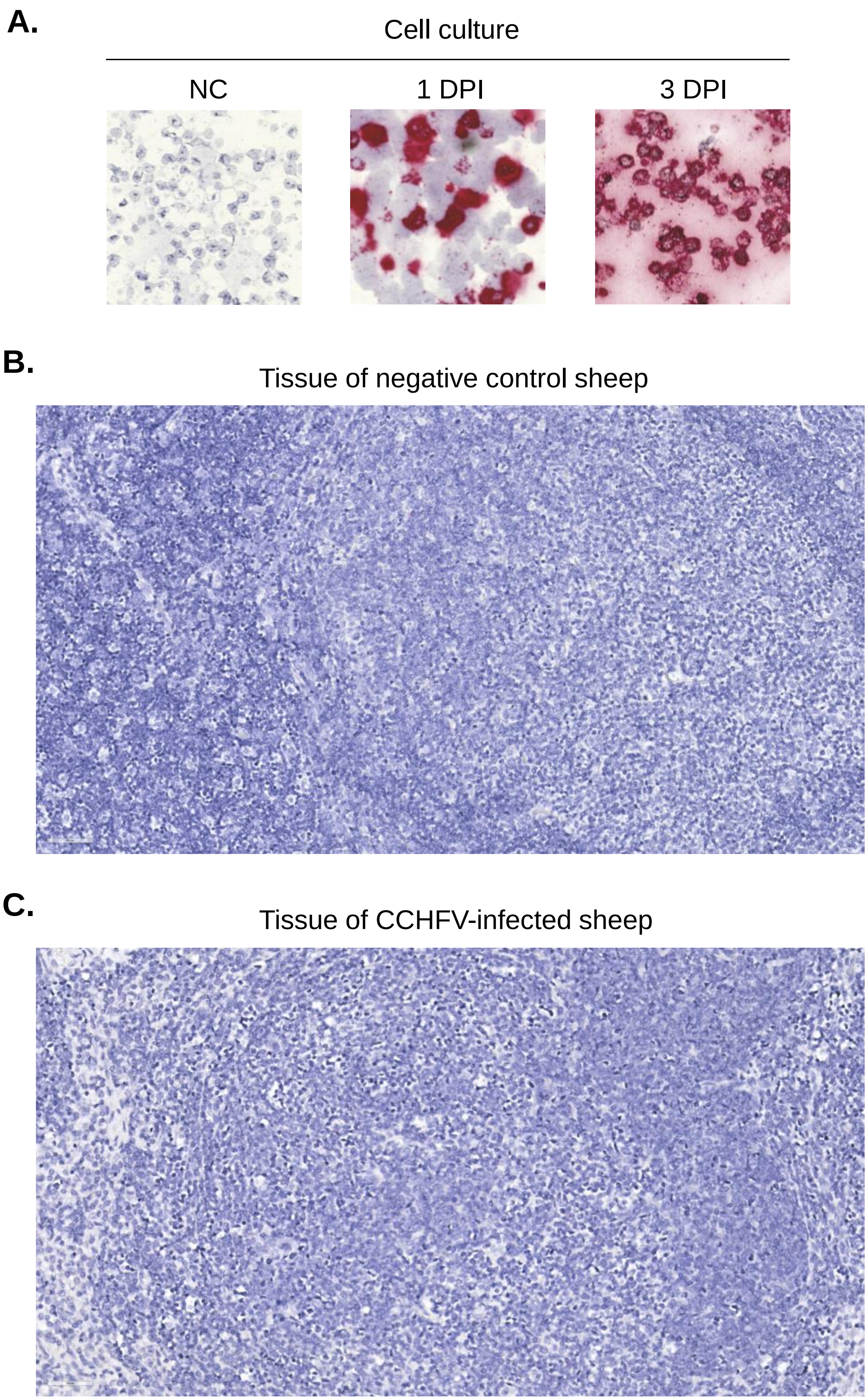

**Appendix Figure 9. Establishment of ISH for positive-sense CCHFV RNA (derived from the M segment). A.** Images of control staining using cell pellets from CCHFV cell cultures. NC, negative control cell culture without CCHFV. 1 DPI and 3 DPI, positive control cell cultures with CCHFV on 1 and 3 day(s) post-infection, respectively. **B.** Image of CCHFV-negative control staining using a lymph node tissue from a sheep infected with foot-and-mouth disease virus. **C.** Image representing negative staining in tissues from CCHFV-infected sheep. Image representing positive staining is shown in Figure 2B. A summary of results from multiple sheep tissues are provided in Appendix Table 3. Images were taken at 40× magnification and scaled down in this figure to 12% relative to original size.

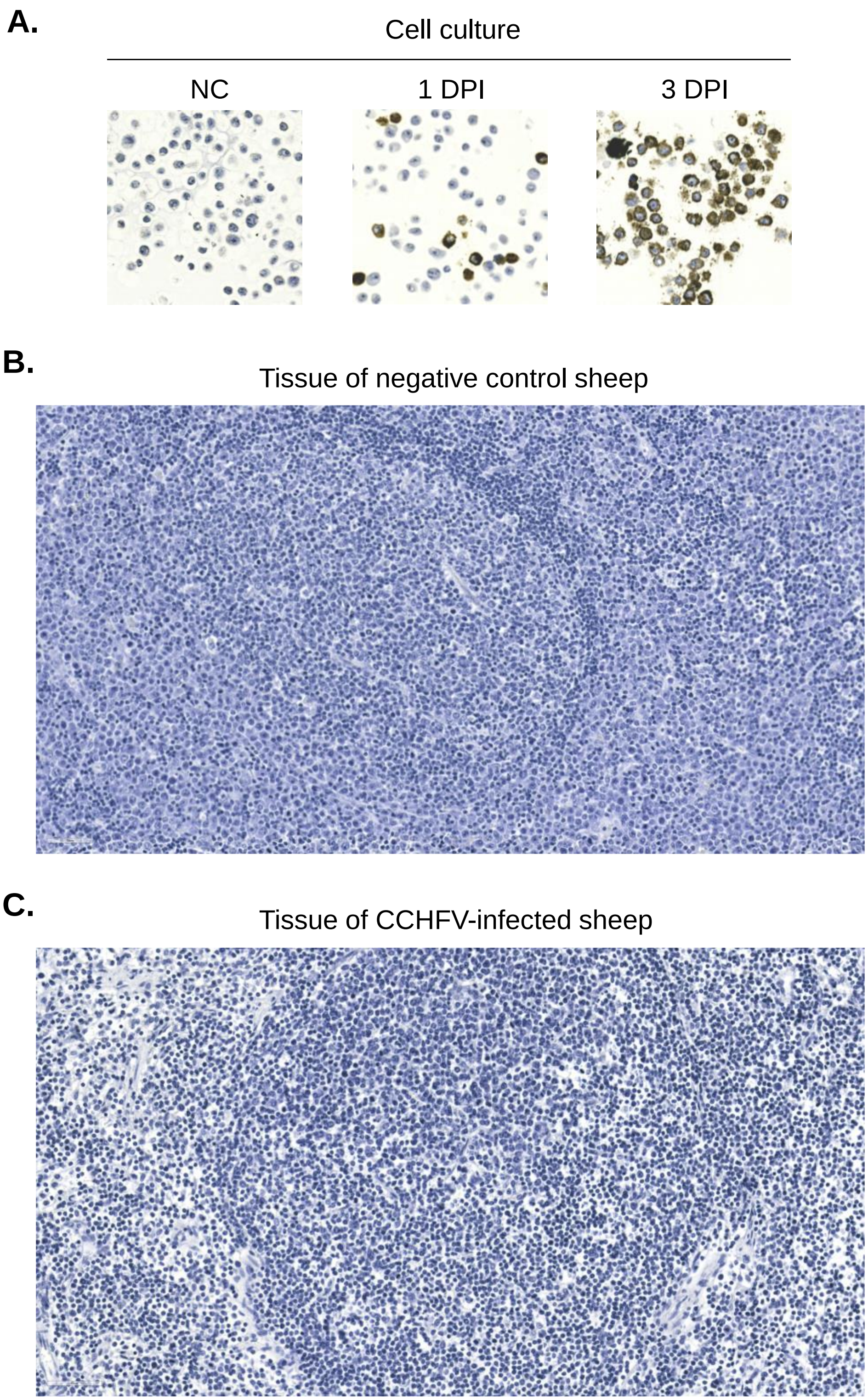

**Appendix Figure 10. Establishment of immunohistochemistry (IHC) for CCHFV N protein. A.** Images of control staining using cell pellets from CCHFV cell cultures. NC, negative control cell culture without CCHFV. 1 DPI and 3 DPI, positive control cell cultures with CCHFV on 1 and 3 day(s) post-infection, respectively. **B.** Image of CCHFV-negative control staining using a lymph node tissue from a sheep infected with foot-and-mouth disease virus. **C.** Image representing staining in tissues from CCHFV-infected sheep, which was generally negative as shown here, with rare exception of positive staining as shown in Figure 2C. A summary of results from multiple sheep tissues are provided in Appendix Table 3. Images were taken at 40× magnification and scaled down in this figure to 12% relative to original size.

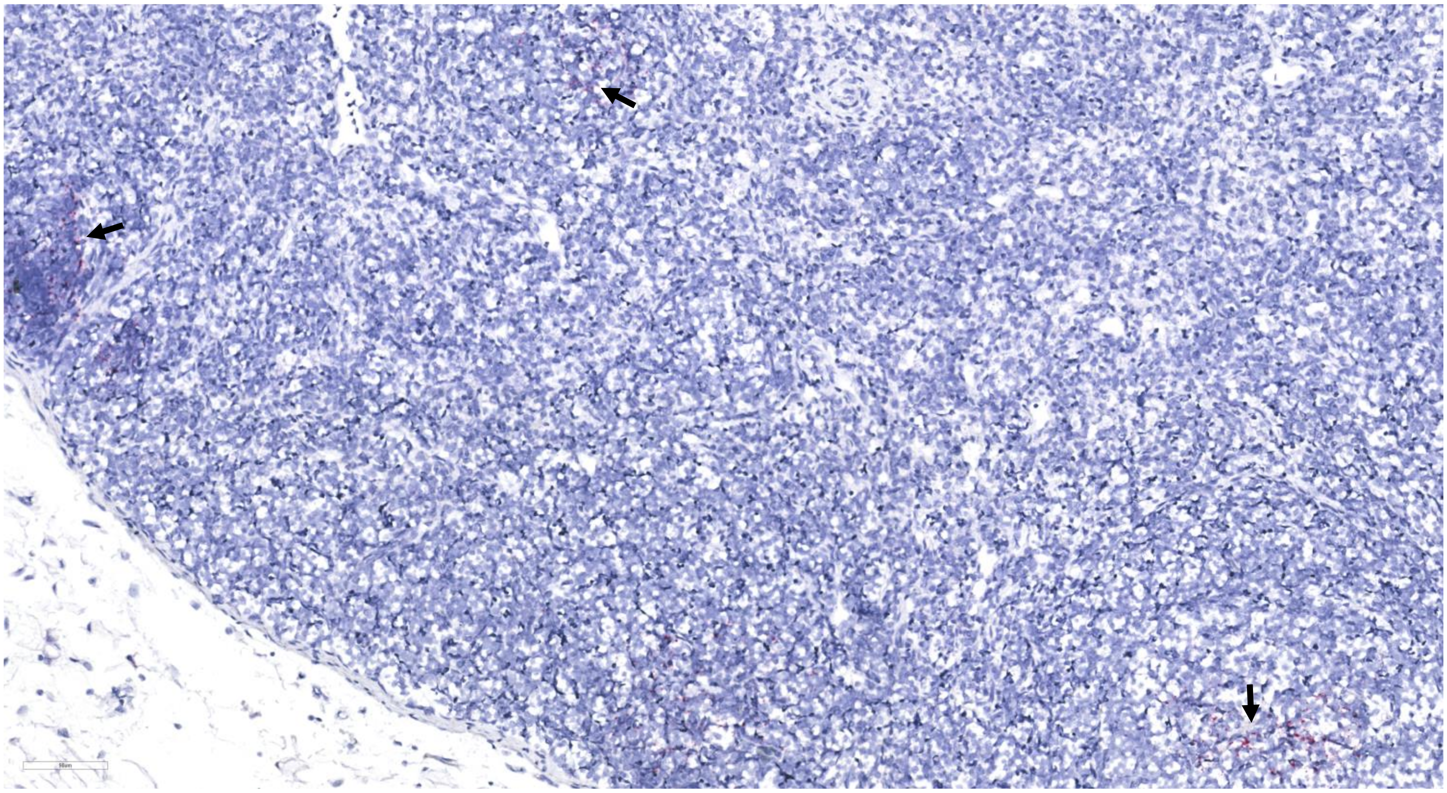

**Appendix Figure 11. Image of ISH for negative-sense CCHFV RNA (M segment) displaying a larger area of tissue section.** Image was taken at 20× magnification and scaled down in this figure to 19% relative to original size. Arrows point to areas of positive staining.

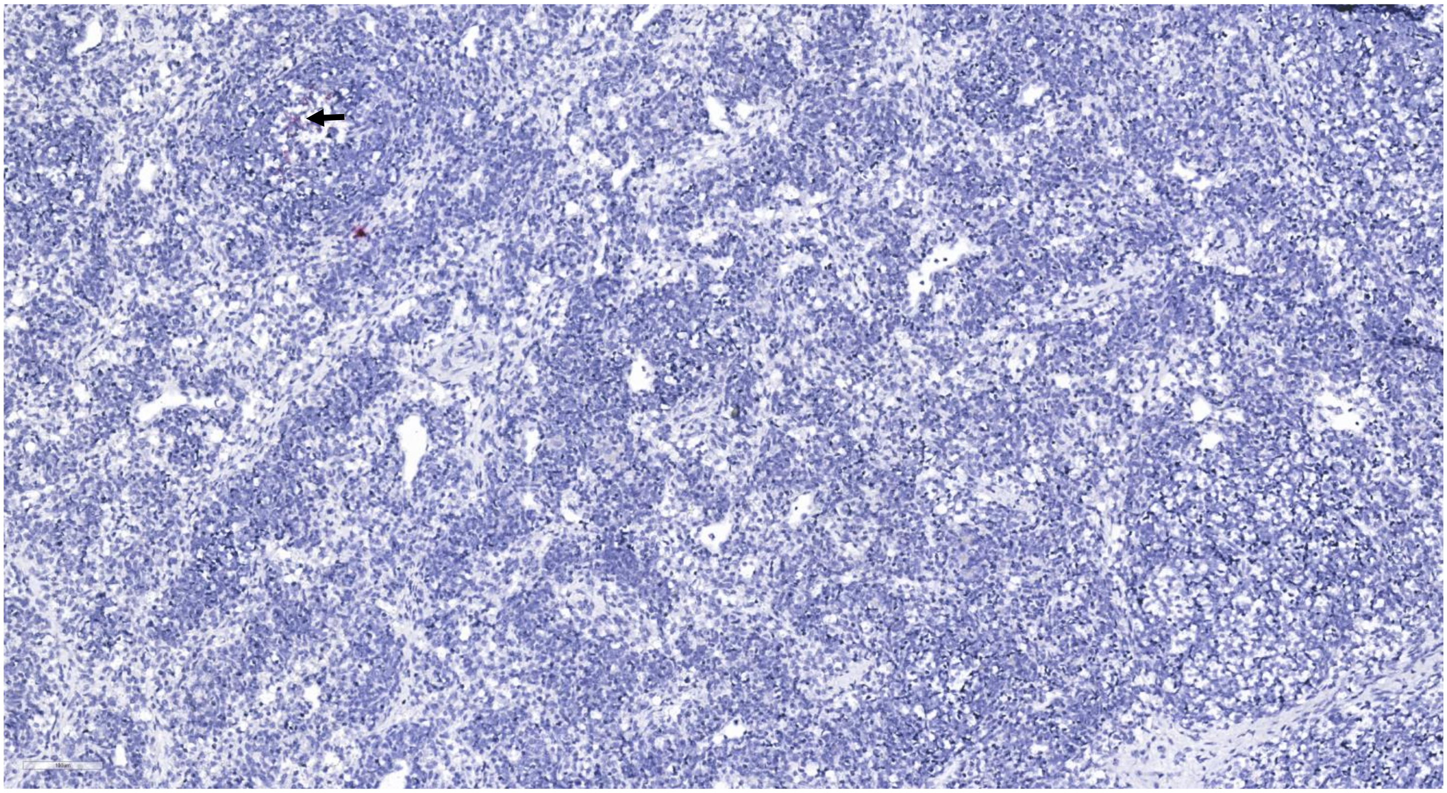

**Appendix Figure 12. Image of ISH for positive-sense CCHFV RNA (M segment) displaying a larger area of tissue section.** Image was taken at 20× magnification and scaled down in this figure to 19% relative to original size. Arrow points to area of positive staining.

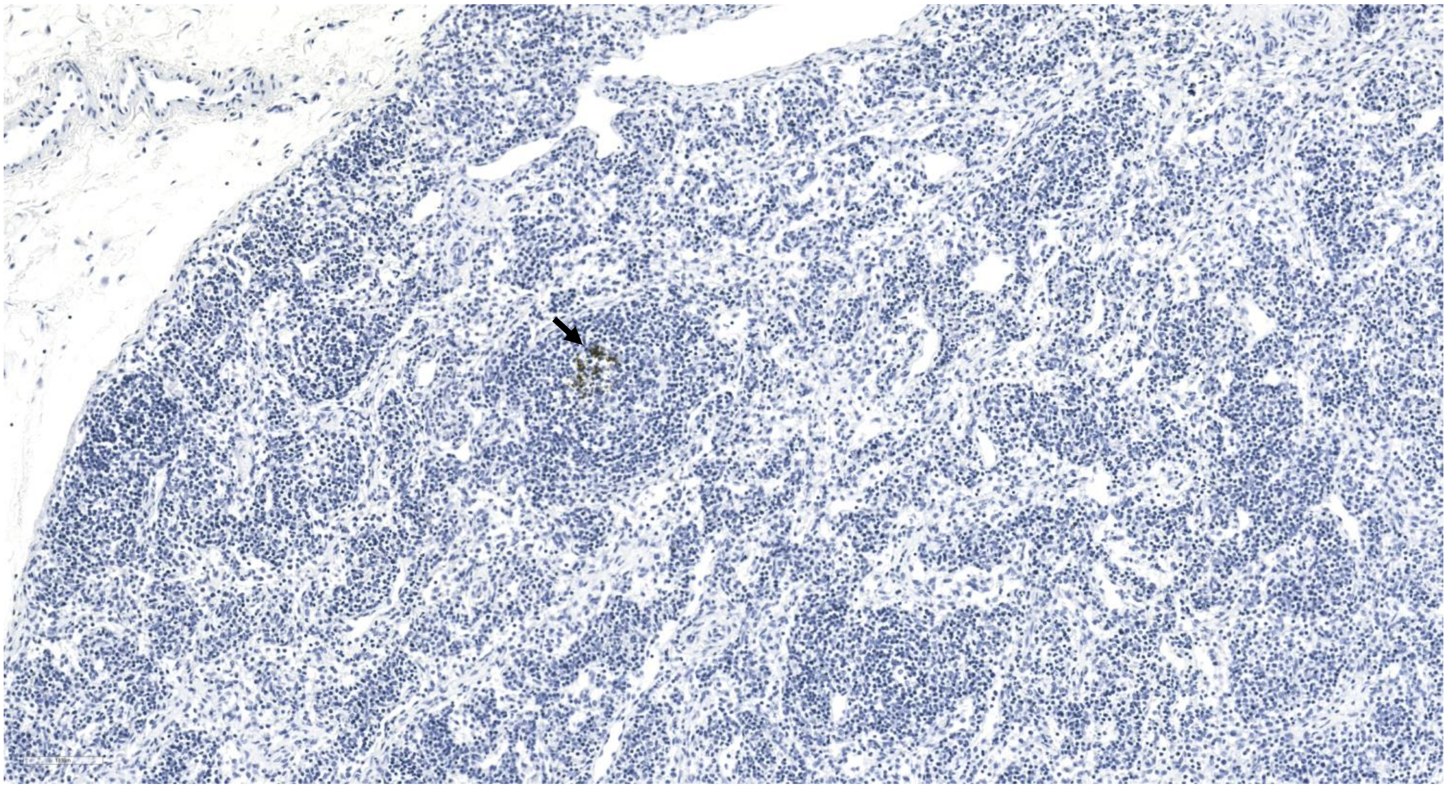

**Appendix Figure 13. Image of IHC for CCHFV N protein displaying a larger area of tissue section.** Image was taken at 20× magnification and scaled down in this figure to 19% relative to original size. Arrow points to area of positive staining.

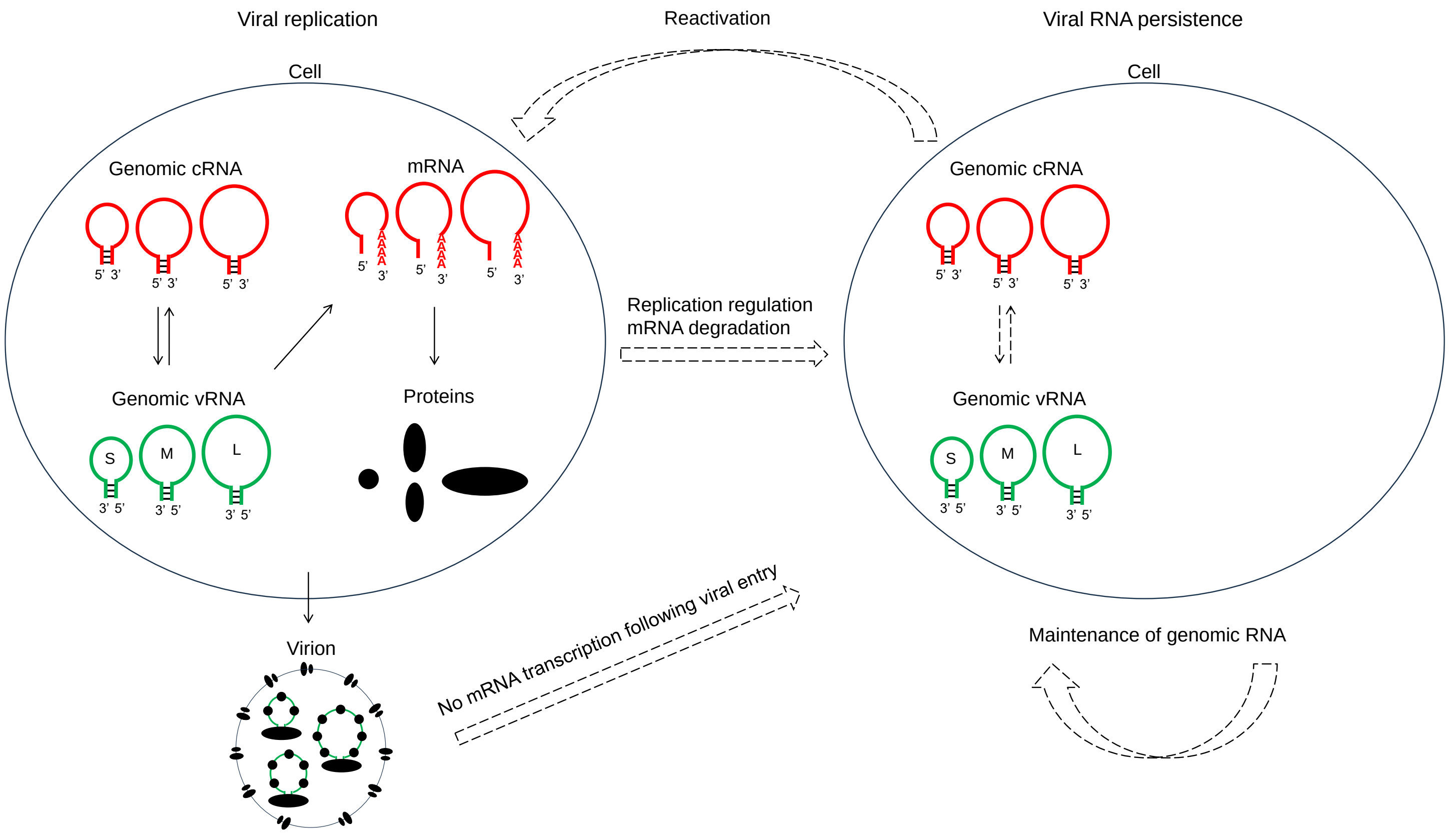

**Appendix Figure 14. Hypothetical model of CCHFV RNA persistence in tissues.** Several pathways are proposed that may potentially contribute to viral RNA persistence. The core reservoir harbors genomic vRNA and cRNA intracellularly and lacks extensive mRNA transcription and virion production. Sporadic reactivation of viral replication, however, could occur in response to immune control relaxation and external stimuli. Viral RNA persistence could originate through two possible routes, depending on the type and antiviral state of the host cell. During a recent viral replication, the host cell may proceed to a relatively quiescent state in which virion production is inhibited and viral mRNA is degraded while genomic vRNA and cRNA are retained. Alternatively, viral entry into a new host cell may be followed only by genomic replication, with mRNA transcription inhibited and no virion produced. Concerning the maintenance of genomic vRNA and cRNA, they may remain in a static state in long-lived cells or replicate using each other as a template and be passed on to new cells through cell division or cell-to-cell transfer of viral RNPs.
